# Unraveling latent cognitive, metacognitive, strategic, and affective processes underlying children’s problem-solving using Bayesian cognitive modeling

**DOI:** 10.1101/2025.01.29.635409

**Authors:** Percy K Mistry, Hyesang Chang, Dawlat El-Said, Vinod Menon

## Abstract

Children exhibit remarkable variability in their mathematical problem-solving abilities, yet the cognitive, metacognitive and affective mechanisms underlying these individual differences remain poorly understood. We developed a novel Bayesian model of arithmetic problem-solving (BMAPS) to uncover the latent processes governing children’s arithmetic strategy choice and efficiency. BMAPS inferred cognitive parameters related to strategy execution and metacognitive parameters related to strategy selection, revealing key mechanisms of adaptive problem solving. BMAPS parameters collectively explained individual differences in problem-solving performance, predicted longitudinal gains in arithmetic fluency and mathematical reasoning, and mediated the effects of anxiety and attitudes on performance. Clustering analyses using BMAPS parameters revealed distinct profiles of strategy use, metacognitive efficiency, and developmental change. By quantifying the fine-grained dynamics of strategy selection and execution and their relation to affective factors and academic outcomes, BMAPS provides new insights into the cognitive and metacognitive underpinnings of children’s mathematical learning. This work advances powerful computational methods for uncovering latent mechanisms of complex cognition in children.

## Introduction

Mathematical knowledge is a critical foundation for success in modern society, yet children show vast individual differences in problem-solving abilities that are not well understood ^1–9^. Identifying the cognitive mechanisms that underlie this variability is essential for designing educational interventions that support the learning needs of all students. While substantial research has examined the role of numerical processing, memory retrieval, cognitive control, and metacognition in math performance^10–14^, a key challenge lies in disentangling how these multiple processes interact to shape problem solving dynamics and skill development. Importantly, students’ problem-solving strategies have been found to vary within individuals, across problems, and over time, necessitating a trial-level approach to elucidate the complex dynamics of strategy adaptivity and efficiency.

One important factor that has received increasing attention is the diversity of strategies that children employ when solving arithmetic problems^15–20^. Even for a simple addition problem like 8 + 5, a student might retrieve the answer from memory, decompose the addends into smaller parts (e.g., 8 + 2 + 3), or count up from the larger number. These strategies have been shown to vary not only across individuals but also within individuals across problems and over time^15,17,18,21–27^. Such strategic variability is thought to reflect the dynamic interplay of cognitive capacities, metacognitive monitoring, and contextual factors^15,16,18,21–27^.

However, conventional methods for studying strategy use have been limited in their ability to capture these complex dynamics. Most previous studies have relied heavily on aggregate accuracy or response time measures that obscure these fine-grained variations^28,29^. Verbal self-reports of strategies provide a limited and potentially biased window into cognitive processes, especially in younger children^30–34^. Moreover, the vast majority of research has focused on snapshots of performance at a single time point, leaving open questions about how problem-solving strategies evolve with learning and development.

Computational models offer a promising approach for addressing these limitations by formalizing explicit hypotheses about the mechanisms underlying children’s problem solving^18^ . However, previous models have not unpacked the sources of individual differences in children’s problem-solving behavior or the dynamics of strategy selection and execution at a fine-grained trial level. Crucially, previous studies have failed to incorporate metacognitive processes involved in monitoring and controlling one’s own cognitive processes during problem solving^22,35–39^. This includes the ability to accurately assess the effectiveness of different strategies for a given problem (monitoring) and to adaptively select and switch between strategies based on this assessment (control).

To fill this gap, we developed a novel Bayesian model of arithmetic problem-solving and strategy-selection (BMAPS) that makes unsupervised inferences about children’s strategy use and efficiency on a trial-by-trial basis. Our model builds on key insights of previous work ^40^ while introducing several innovations to capture the richness of problem-solving dynamics. First, it specifies distinct cognitive parameters related to the execution of different strategies, including the efficiency of fact retrieval and procedural computation. Second, it includes metacognitive parameters related to the adaptivity and flexibility of strategy selection, such as the time to switch between strategies and the ability to calibrate strategy choices to problem difficulty. Finally, it incorporates hierarchical structures to estimate individual differences in these cognitive and metacognitive components while also identifying global patterns of strategy use and performance.

We leveraged this model to test several hypotheses about the mechanisms underlying children’s arithmetic performance and learning (**Figure 1**). Our first aim was to evaluate whether the cognitive and metacognitive parameters identified by the model could explain unique variance in problem-solving accuracy, response times, and error rates beyond that accounted for by aggregate performance measures. We expected that efficiency of fact retrieval and adaptivity of strategy selection would be key predictors of problem-solving success.

**Figure 1:**
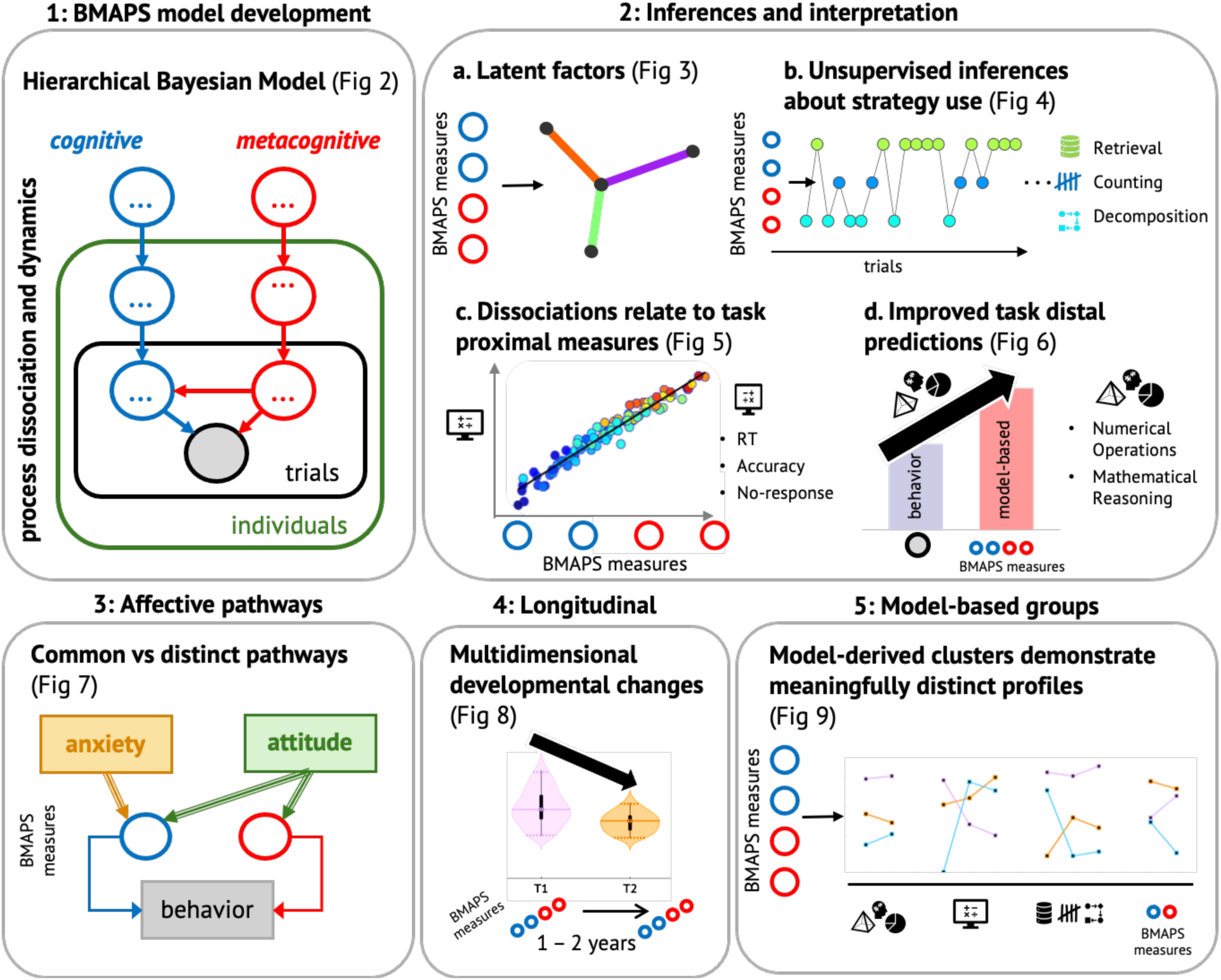
Overview of aims and analyses. Schematic representation of the data analysis approach used in this study to obtain model-derived insights.

Our second aim was to assess the power of BMAPS parameters to predict individual differences in standardized measures of mathematical achievement. We hypothesized that students with higher retrieval efficiency and strategy adaptivity would show advantages not only in arithmetic computation but also in broader assessments of mathematical concepts and problem solving.

This would suggest that the cognitive skills captured by our model are foundational for the development of mathematical competence.

Our third aim was to elucidate the specific pathways by which affective factors like math anxiety and math attitudes influence problem-solving performance. Previous research has linked math anxiety to reduced use of problem-solving strategies^41^ and lower arithmetic fluency^42,43^, but the mechanisms underlying these effects are not clear. Here we test the hypothesis that math anxiety impacts performance by reducing the efficiency of fact retrieval, while positive attitudes promote performance by increasing the adaptivity of strategy selection. Identifying these precise relationships could inform efforts to foster positive academic mindsets and mitigate the deleterious effects of anxiety.

Our fourth aim was to examine how problem-solving strategies change over the course of learning and development. We used longitudinal data to map the trajectories of strategy use and efficiency across one year of schooling, allowing us to identify leading indicators of growth in mathematical competence. We hypothesized that students who showed greater gains in retrieval efficiency and strategy adaptivity would also show larger improvements in problem-solving accuracy and mathematical achievement. Charting these developmental dynamics is critical for understanding how strategic expertise emerges over time and how educational interventions can support this process^44^

To test these hypotheses, we applied BMAPS to a rich dataset of second- and third-grade students (*N* = 105) who completed a battery of arithmetic problem-solving tasks, standardized achievement tests, and attitude surveys at two time points. This approach allowed us to map the fine-grained landscape of children’s problem-solving behavior, identify sources of individual differences in performance and growth, and relate cognitive processes to affective factors and academic outcomes.

In summary, we present a novel computational approach for unpacking the complex dynamics of strategy use and efficiency that underlie children’s mathematical problem solving. Our model extends previous work by providing a unified framework for estimating trial-level variations in cognitive and metacognitive processes, identifying sources of individual differences in performance and growth, and relating problem-solving behavior to broader measures of achievement and affect. We demonstrate the utility of this approach through a series of analyses that test key hypotheses about the mechanisms of children’s strategic development. Our findings highlight the importance of retrieval efficiency and strategy adaptivity as foundational skills for mathematical learning, and suggest promising avenues for educational interventions that target these capacities. More broadly, this work showcases the power of computational methods for advancing our understanding of the cognitive bases of academic achievement and informing evidence-based efforts to support all learners.

## Results

### Bayesian model of arithmetic problem solving (BMAPS)

BMAPS infers strategy choice and execution at a single trial level by implementing a probabilistic mixture model of drift diffusion processes (**Figure 2A**). **Figure 2B** shows a schematic of a sample trial where the default retrieval processes initiates but is not resolved by the switching time, and a probabilistic decision is made at the switching time to discontinue retrieval and initiate counting. On each trial, if the retrieval process is not resolved by the switching time, a probabilistic decision may be taken to still continue with retrieval, or switch to either counting or decomposition strategies. The model allows inferences about switching time, strategy efficiency (drift rate), uncertainty over strategy choice, and probabilistic strategy selection on each trial, governed by hierarchical cognitive and metacognitive processes (see **Methods** for details).

**Figure 2:**
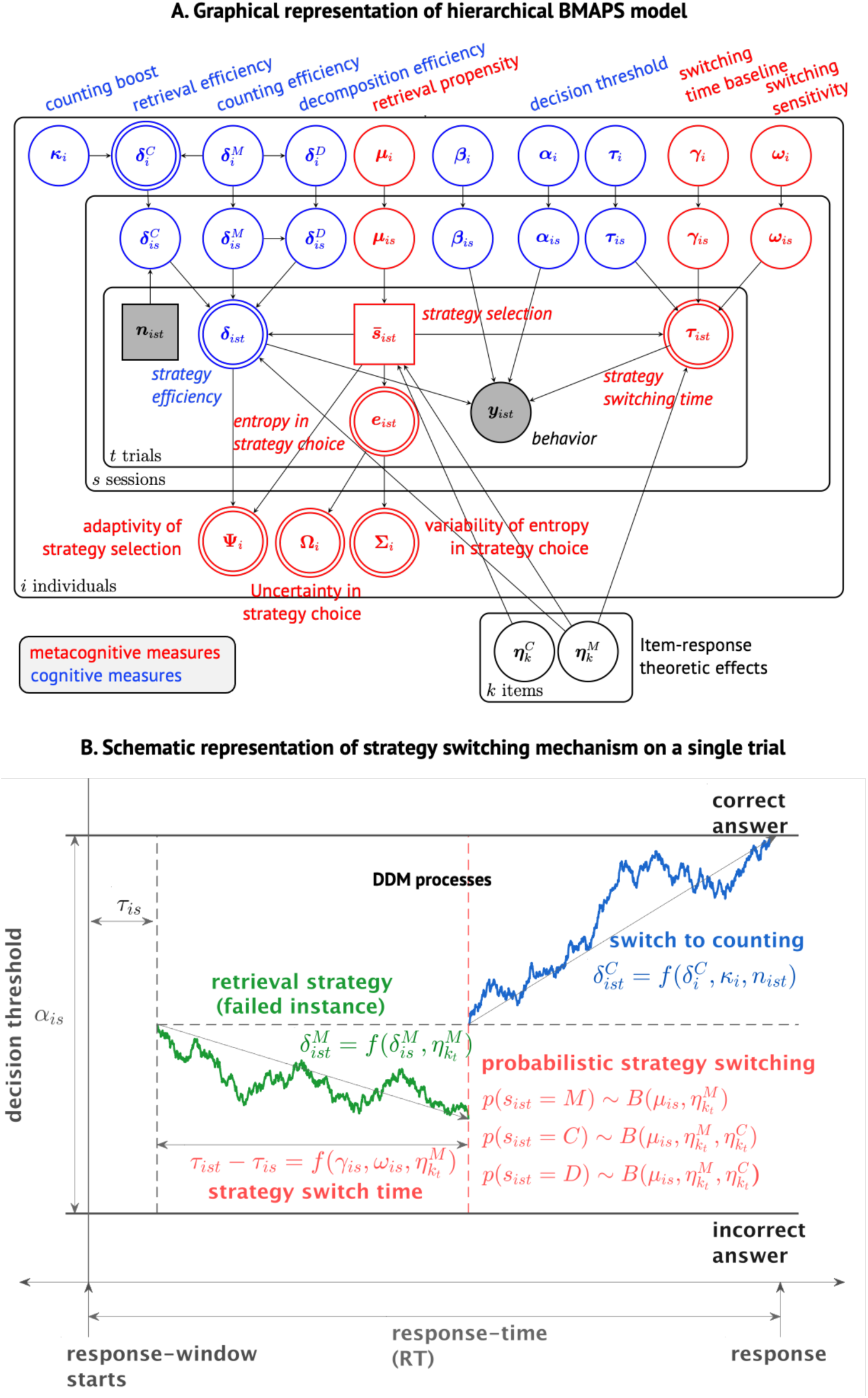
**A. Graphical model showing the hierarchical Bayesian implementation of BMAPS and the underlying latent cognitive and metacognitive measures.** *y* represents the observed choice (accuracy) coded response times. The distribution of choice coded response times is modeled as a mixture of drift diffusion model (DDM). Nodes in circles represent continuous measures, while squares represent discrete values. Shaded nodes are observed values while the remaining are latent variables. Nodes with a single border are probabilistic (inferred values), while those with double borders are deterministic (derived, or calculated). Finally, nodes in red refer to the metacognitive components of BMAPS, and nodes in blue are item-specific (problem-specific) components. **Table 1** provides the description of BMAPS variables. **B. Schematic of model components.** Schematic of a sample trial where the default retrieval processes initiates but is not resolved by the switching time, and a probabilistic decision is made at the switching time to discontinue retrieval (green) and initiate counting (red). The model allows inferences about switching time, strategy efficiency (drift rate), uncertainty over strategy choice, and probabilistic strategy selection on each trial, governed by hierarchical cognitive and metacognitive processes which are characterized by BMAPS parameters.

**Table 1:**
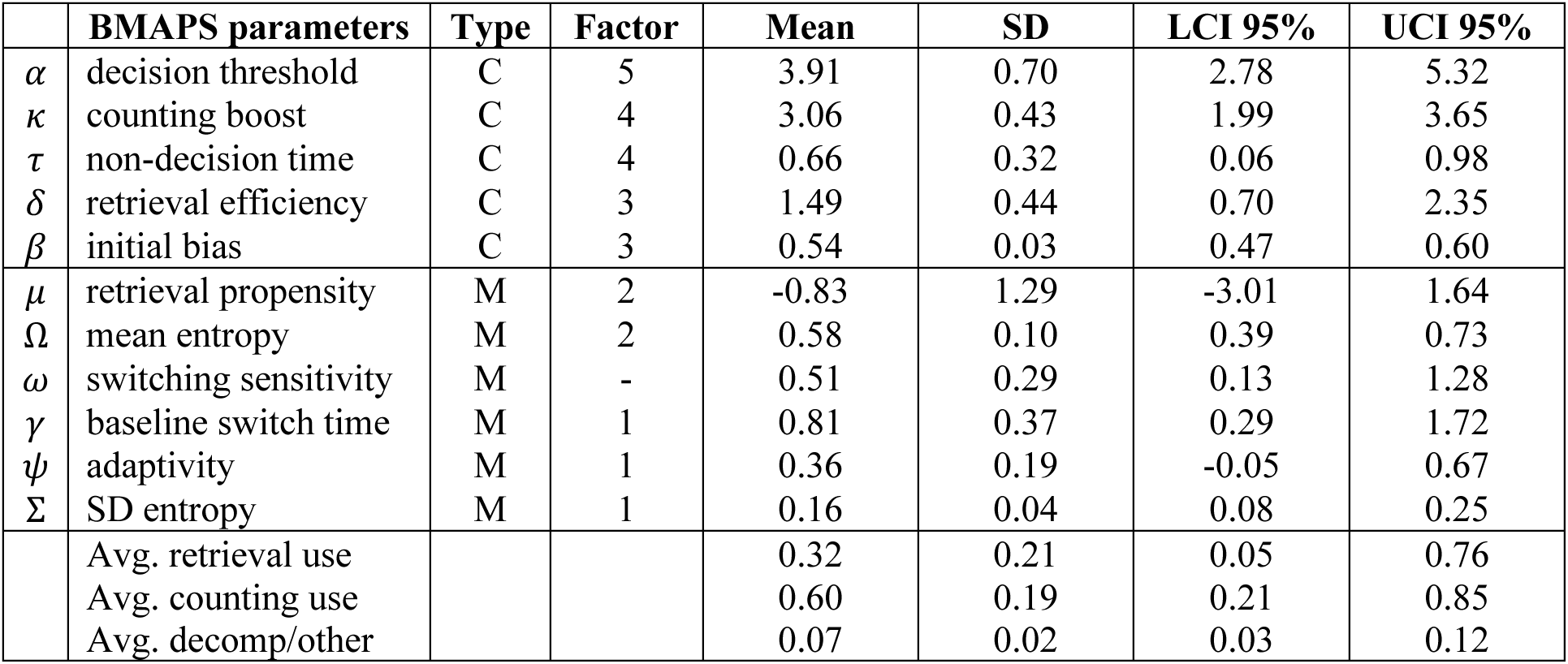
Summary of key BMAPS individual-level model parameters. The table shows the mean, SD, and 95%CI for model inferred parameters across the 105 participants (excluding outliers; see Methods). Type ‘C’ parameters are cognitive parameters related to strategy execution and performance, and are parameters typically associated with standard drift diffusion models. Type ‘M’ parameters are metacognitive parameters related to strategy switching and selection, and are novel measures introduced in this model that hierarchically govern the switching and modulation of the drift diffusion model processes. The factor column shows the factor of the 5-factor model that the parameters are most strongly associated with.

|  | **BMAPS parameters** | **Type** | **Factor** | **Mean** | **SD** | **LCI 95%** | **UCI 95%** |
| --- | --- | --- | --- | --- | --- | --- | --- |
| $\alpha$ | decision threshold | C | 5 | 3.91 | 0.70 | 2.78 | 5.32 |
| $\kappa$ | counting boost | C | 4 | 3.06 | 0.43 | 1.99 | 3.65 |
| $\tau$ | non-decision time | C | 4 | 0.66 | 0.32 | 0.06 | 0.98 |
| $\delta$ | retrieval efficiency | C | 3 | 1.49 | 0.44 | 0.70 | 2.35 |
| $\beta$ | initial bias | C | 3 | 0.54 | 0.03 | 0.47 | 0.60 |
| $\mu$ | retrieval propensity | M | 2 | -0.83 | 1.29 | -3.01 | 1.64 |
| $\Omega$ | mean entropy | M | 2 | 0.58 | 0.10 | 0.39 | 0.73 |
| $\omega$ | switching sensitivity | M | - | 0.51 | 0.29 | 0.13 | 1.28 |
| $\gamma$ | baseline switch time | M | 1 | 0.81 | 0.37 | 0.29 | 1.72 |
| $\psi$ | adaptivity | M | 1 | 0.36 | 0.19 | -0.05 | 0.67 |
| $\Sigma$ | SD entropy | M | 1 | 0.16 | 0.04 | 0.08 | 0.25 |
|  | Avg. retrieval use |  |  | 0.32 | 0.21 | 0.05 | 0.76 |
|  | Avg. counting use |  |  | 0.60 | 0.19 | 0.21 | 0.85 |
|  | Avg. decomp/other |  |  | 0.07 | 0.02 | 0.03 | 0.12 |

### BMAPS provides robust inference of strategy use

Verbal self-reports have well-documented limitations for assessing children’s problem-solving strategies: they can be influenced by task instructions and experimenter expectations and are particularly unreliable in young children with limited metacognitive awareness^31–33^. They are difficult to collect online during a task, without interfering with the cognitive processes. We therefore evaluated whether BMAPS could provide robust strategy assessment of tasks completed within the fMRI scanner without relying on self-reports.

Given that strategy use naturally varies across problems and sessions even within individuals and self-reports may include a degree of noise, we first established empirical benchmarks of within-individual and between-individual variability in independently collected self-reported strategies (**Figure S1, SI Table S1**). Within-individual comparisons across different problem sets showed moderate consistency (Kendall’s tau: 0.283; MSE: 0.062). In contrast, between-individual comparisons showed poor consistency (Kendall’s tau: -0.229; MSE: 0.194), indicating distinct individual differences in strategy preferences. We then validated BMAPS-inferred strategy use against self-reports and compared the resulting variability to the empirical benchmarks established above. We found that that BMAPS strategy inferences showed within-individual consistency comparable to self-reports, while successfully capturing individual differences in strategy preferences (Kendall’s tau: 0.193, MSE: 0.094). These results demonstrate that our modeling approach can reliably assess strategy use without the limitations and biases of self-reports. See **Methods** and **SI** for detailed validation procedures.

### BMAPS outperforms behavioral measures in predicting self-reported strategy use

To further validate our modeling approach, we compared its ability to predict self-reported strategy use against traditional model-free methods based on behavioral measures (accuracy, reaction time) and neuropsychological assessments (working memory, IQ). Using cross-validated regression analyses, our model-based predictions of retrieval strategy use significantly outperformed predictions from model-free methods (mean Kendall’s tau: 0.231 vs. 0.146, effect size = 1.923, p < 0.0001; mean MSE: 0.067 vs. 0.076, effect size = -1.963, p < 0.0001; **Supplementary Figure S1**). These findings demonstrate that BMPAS captures meaningful aspects of strategy use that are not readily apparent from observable behavioral measures alone.

### Latent BMAPS cognitive and metacognitive measures capture multidimensional mechanisms of adaptive problem solving

BMAPS identified latent parameters related to problem-solving execution and strategy selection. Execution parameters included retrieval efficiency δ (speed and accuracy of fact retrieval), counting boost κ (efficiency of counting strategies), and decision threshold α (evidence required for a decision). Strategy selection parameters included retrieval propensity μ (likelihood of using retrieval), baseline switching time γ (time to switch between strategies), and switching sensitivity ω (modulation of switching time based on item difficulty). We also derived measures of adaptivity ψ (optimality of retrieval strategy selection), mean entropy Ω (average uncertainty in strategy selection), and standard deviation of entropy Σ (variability in strategy uncertainty across trials) from trial-level parameters (see **Methods** for details). Descriptive statistics of model parameters and strategy use proportions are shown in **Table 1**. Correlations between parameters (**Figure 3A, SI Table S2**) revealed that baseline switching time γ was the only metacognitive parameter significantly correlated with cognitive parameters.

**Figure 3:**
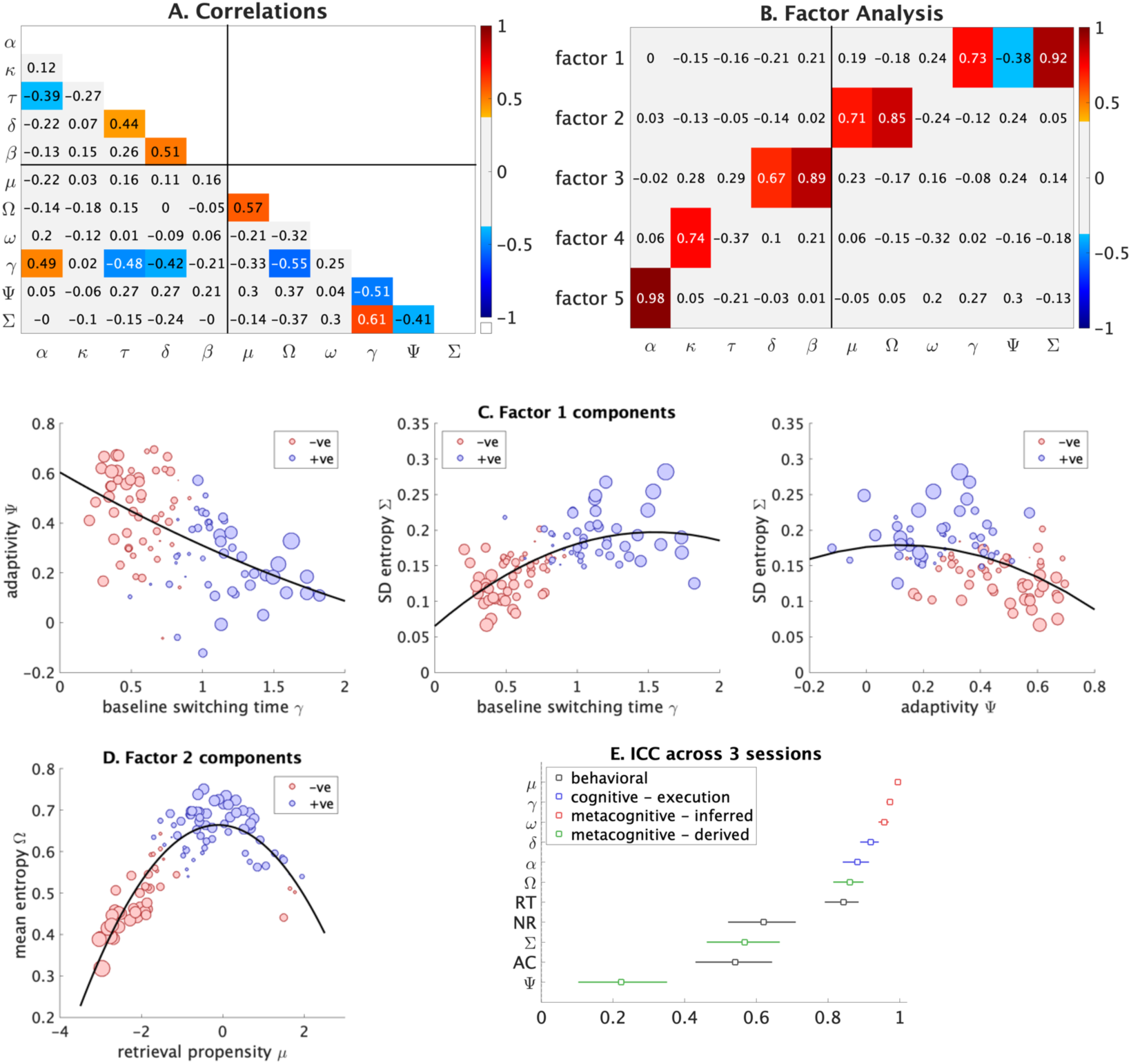
**A. Correlations between BMAPS parameters.** Significant correlations (p <=0.05) after adjusting for multiple comparisons are presented in a range of colors based on effect size. Values represent Pearson’s *r*. Significant positive correlations are shown in orange/red cells and significant negative correlations are shown in light/dark blue cells. **B. Factor analysis of model parameters.** BMAPS parameter loadings on five factors. Significant positive loadings are shown in orange/red cells and significant negative loadings are shown in light/dark blue cells. Metacognitive measures load on to factors 1 and 2, while cognitive measures load on to factors 3, 4, and 5. **C. Relationships between key component measures of factor 1.** The plots show the pairwise associations between baseline switching time, adaptivity, and SD of entropy. The size of the dots represent the absolute value of factor scores and the color of dots correspond to negative (red) or positive (blue) factor scores. **D. Relationships between key component measures of factor 2.** The plot shows the associations between retrieval propensity and mean entropy. The size of the dots represent the absolute value of factor scores and the color of dots correspond to negative (red) or positive (blue) factor scores. **E. Intra-class correlations across 3 sessions.** The figure shows ICC values measured across 3 sessions for behavioral (reaction time [RT], accuracy [AC], and no-response rate [NR]) as well as key BMAPS cognitive and metacognitive parameters. The model was fit hierarchically with individual level priors and inferences at a session and individual level. Metacognitive inferred measures are based on the statistical inferences made by fitting the model to data. Metacognitive derived measures are based on a second level analysis correlating or calculating values across trial-level inferences within each individual. See **Table 1** for description of BMAPS variables.

Factor analysis showed that 2-factor (*χ*^2^ = 133, p < 0.0001), 3-factor (*χ*^2^ = 83, p < 0.0001), and 4- factor (*χ*^2^ = 42, p = 0.007) models rejected the null hypothesis of 2, 3, 4 factors respectively. A 5-factor model (*χ*^2^ = 16.8, p = 0.079) failed to reject the hypothesis of 5-factors (**Figure 3B**). The 5-factor model included three cognitive factors related to decision threshold, retrieval efficiency, and counting boost; and two metacognitive factors capturing (mal)adaptivity (high switching time, low adaptivity, high entropy variability) and retrieval propensity (high retrieval use, high mean entropy) (**Figures 3C-D**).

Critically, model parameters dissociated variability in observable aspects of problem-solving performance (**Figure 3, SI Tables S3-4**). Reaction time was related to decision threshold, retrieval efficiency, retrieval propensity, switching time, and initial response bias. Accuracy was associated with retrieval efficiency, adaptivity, and bias. No-response rate was associated with decision threshold, retrieval efficiency, switching time, switching sensitivity, mean entropy, and bias. Furthermore, metacognitive parameters of retrieval propensity, baseline switching time, and switching sensitivity showed high levels of between-session stability (ICC > 0.95) compared to behavioral measures (ICCs 0.54 - 0.84), suggesting their potential utility as stable cognitive markers (**Figure 3E**).

Finally, we assessed all BMAPS measures and factor scores for sex differences and found no differences on any of the measures (**Supplementary Table S5).**

### BMAPS captures strategy- and item-dependent performance differences

To further validate our model, we assessed performance differences between model-inferred strategies and items of varying difficulty. As expected, reaction times increased from retrieval to counting to decomposition strategies, with larger differences between easy (sum ≤ 10) and hard (sum > 10) problems for counting (**Table 2**). Error rates were lowest for counting and highest for decomposition strategies. The distributions of unsupervised model-inferred strategy reaction times aligned well with benchmarks derived from a supervised learning model^27^ (**Figure 4A**), with the RT at which the likelihood of inferring retrieval versus non-retrieval strategies switched, being about 2.6-2.8 seconds, similar to that inferred by conventional supervised strategy classification methods.

**Figure 4.**
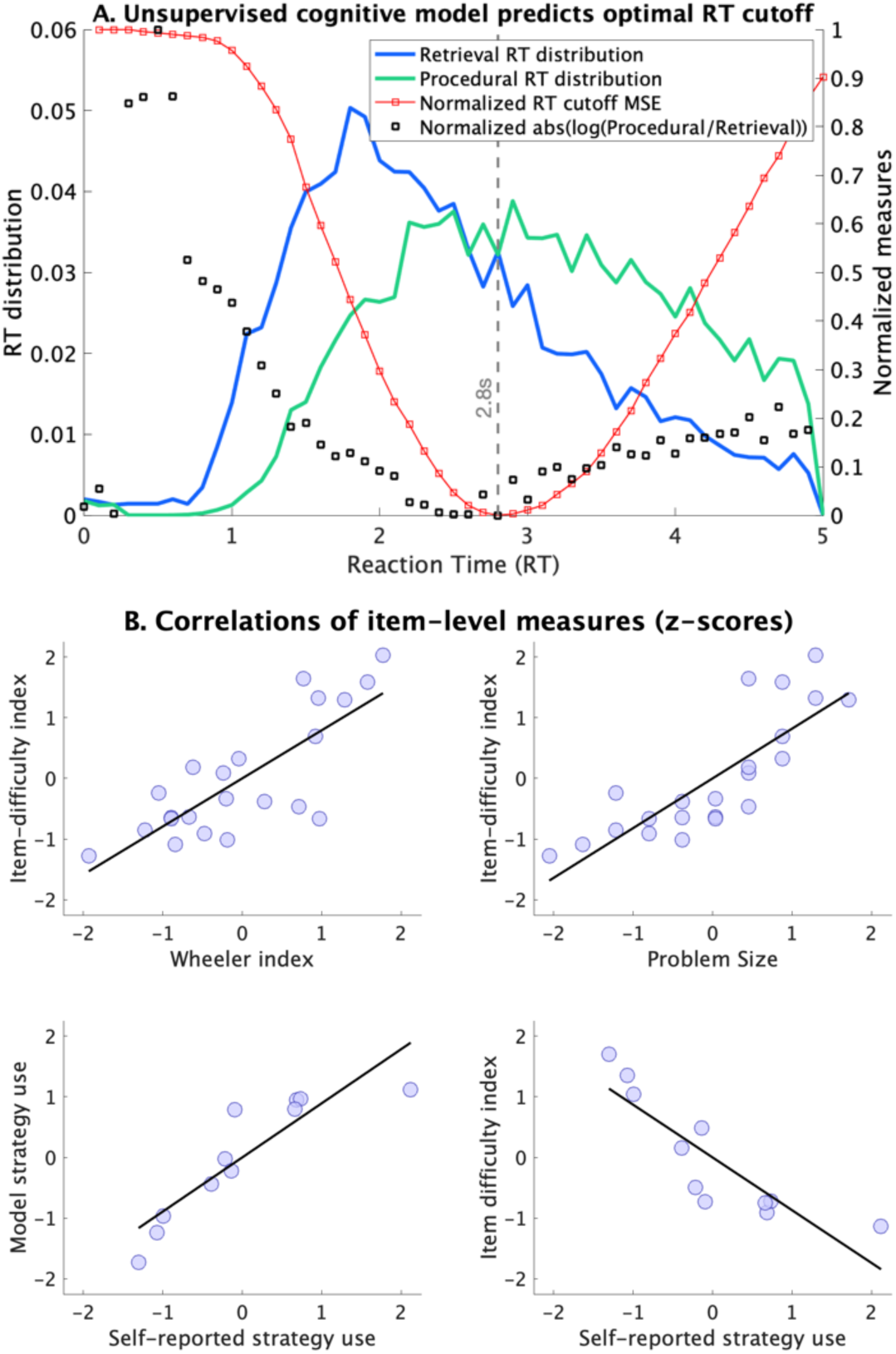
**A. Distribution of reaction times (RTs) for BMAPS inferred retrieval and non-retrieval strategies.** The RT at which the likelihood of observing retrieval vs non-retrieval switches is ∼ 2.6-2.8 seconds, based on the unsupervised cognitive model. To validate the RT distributions of the different strategies in the unsupervised model, a supervised model was used to infer the most optimal RT cut-off such that classifying responses shorter than that RT as retrieval, and longer as non-retrieval would provide the highest rate of correct strategy classification. The optimal RT cut-off from the supervised model was identical to the RT at which likelihoods switched between retrieval and non-retrieval strategy in the unsupervised model. The blue and green lines show the distribution of RTs for retrieval and non-retrieval strategies respectively, and the black dots show the normalized absolute log-ratio of these strategies, with the lowest value indicating the point at which both strategies are equally likely, based on the unsupervised model. The red line shows the normalized mean square error while predicting self-reported strategy use based on the assumption of a best-fitting RT based cut-off for whether or not a particular trial is solved by retrieval. **B. Construct validity of model inferred strategies.** Significant correlations were observed between unsupervised model inferred item-difficulty index (which influences both the probability of using retrieval as well as the efficiency of retrieval execution) and Wheeler index, problem-size, and self- reported strategy. Significant correlation was observed between model-inferred strategy use and self- reported strategy use for the same subset of items included in both in-scanner and out-of-scanner tasks.

**Table 2:** Summary of strategy-specific behavioral characterization. The table shows the mean reaction time, answer error rate, no-response rate, and overall error rate for easy (answer <=10) and hard (answer >10) problem types, separately based on whether the specific trial level strategy choices were classified as retrieval, counting, or decomposition.

|  | **Type** | **Retrieval** | **Counting** | **Other** |
| --- | --- | --- | --- | --- |
| Mean Reaction Time (RT) | easy | 2.52s | 2.91s | 3.19s |
|  | hard | 2.72s | 3.36s | 3.47s |
| Accuracy on answered questions (AC) | easy | 86.8% | 94.7% | 67.6% |
|  | hard | 77.6% | 88.4% | 67.2% |
| No-response rate (NR) | easy | 10.8% | 10.2% | 23.7% |
|  | hard | 16.0% | 23.1% | 28.3% |

Hierarchical clustering of model-inferred item difficulty estimates (**Supplementary Figure S2**) led to a clear separation of easy and hard problems, and these estimates correlated strongly with problem size (r = 0.82) and a well-established difficulty index^45^ (**Figure 4B**; r = 0.79). Moreover, model-inferred item-level strategies were highly consistent with participants’ self-reported strategies on the subset of common problems (**Figure 4B**; r = 0.89). These findings demonstrate the model’s validity in capturing meaningful strategy- and item-related differences in performance.

### BMAPS parameters collectively explain task-proximal arithmetic performance

To assess whether model parameters could collectively explain individual differences in arithmetic performance, we conducted canonical correlation analyses (CCA) with accuracy, reaction time, and no-response rate as outcome variables. CCA with the 5 factors (**Figure 5A**) revealed 3 significant canonical variates (r = 0.92, p<0.0001; r = 0.8, p<0.0001; r = 0.44, p = <0.0001), each representing a distinct dimension of covariation between the factors and performance measures. All 5 factors were associated with behavioral performance, and were associated with unique canonical variables.

**Figure 5.**
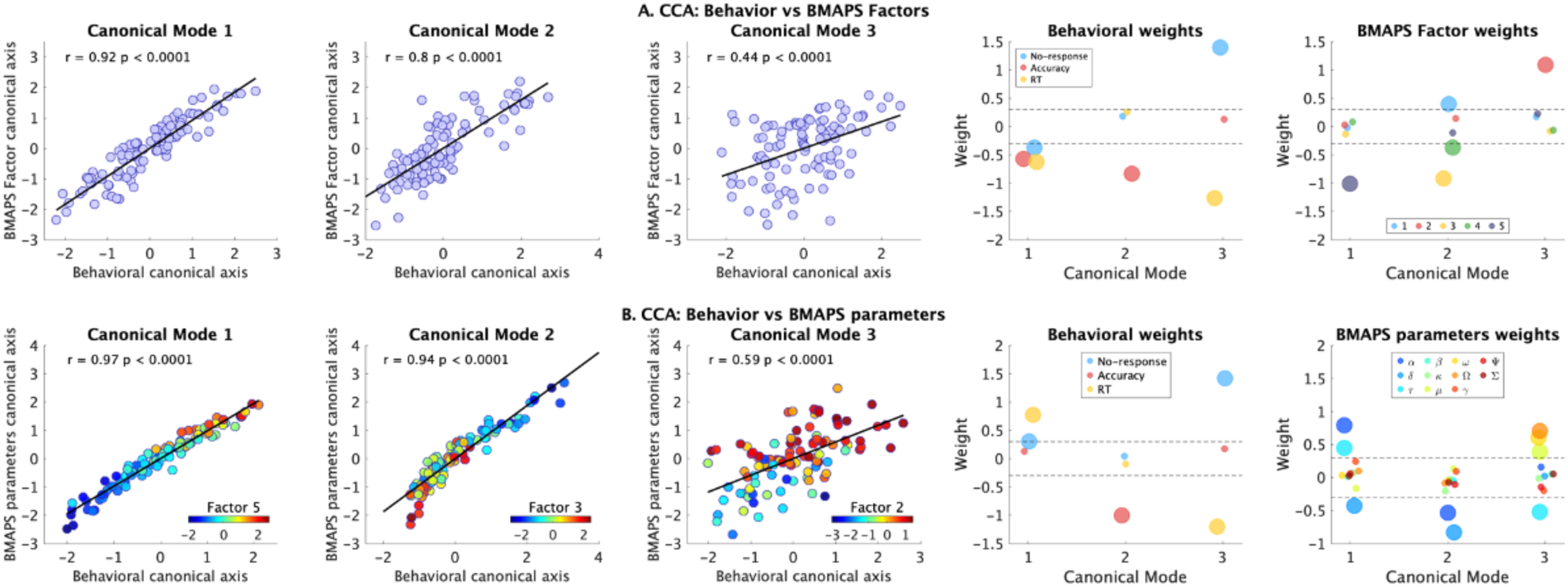
**A. Canonical correlation analyses (CCA) between BMAPS derived factor scores and behavior.** See Figure 3 for BMAPS parameter loadings on five factors. The weights of behavioral and factor variables for each of three significant canonical modes are shown. Weights with absolute value greater 0.3 (dotted lines) are shown in larger circles and those with absolute value less than 0.3 are shown in smaller circles. **B. CCA between BMAPS-inferred measures and behavior.** See **Table 1** for description of BMAPS variables. The weights of behavioral and factor variables for each of three significant canonical modes are shown. Weights with absolute value greater 0.3 (dotted lines) are shown in larger circles and those with absolute value less than 0.3 are shown in smaller circles. The three significant canonical variables showed significant correlations with distinct factor scores (factors 5, 3, 2 respectively). The range of colors in the three plots on the left represents factor scores.

CCA with the 11 original model parameters (**Figure 5B**) similarly yielded 3 significant canonical variates (r = 0.97, p<0.0001; r = 0.94, p<0.0001; r = 0.59, p = <0.0001). These results suggest that our model parameters capture the key dimensions of individual differences in arithmetic problem-solving performance. Importantly, the three significant canonical variates based on BMAPS and behavioral measures show strong correlations with the factor scores (**Figure 5B**).

### BMAPS parameters predict individual differences in broad task-distal mathematical achievement

To test the predictive utility of our model for broader mathematical competencies, we examined correlations between model parameters and standardized measures of arithmetic fluency (Numerical Operations) and mathematical reasoning (Mathematical Reasoning) from the WIAT-II battery. Both measures showed significant correlations (**Figure 6A**) with metacognitive indices of adaptivity, baseline switching time, and SD of strategy selection entropy (all |r| ranging from 0.28 to 0.41, p<= 0.01; **SI Table S6**). Notably, all three of these BMAPS measures load on to the first factor.

**Figure 6.**
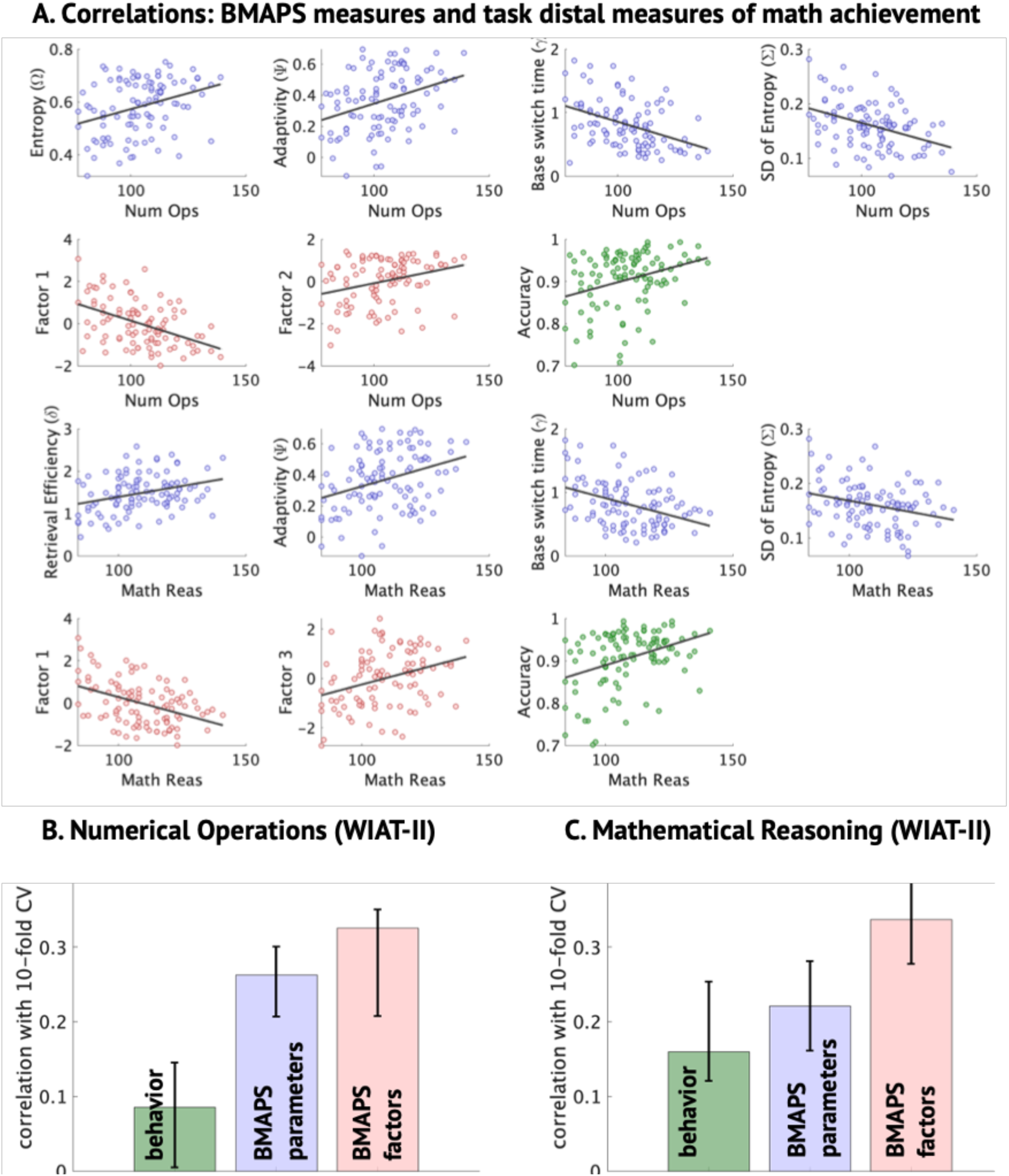
**A. Correlations between math achievement and behavioral and BMAPS measures**. Children’s math achievement was assessed by the Wechsler Individual Achievement Test, Second Edition (WIAT-II) Numerical Operations and Mathematical Reasoning subtests. Mathematical achievement was significantly correlated with behavioral measures (green; higher accuracy on the arithmetic tasks), BMAPS measures (blue; higher entropy, adaptivity, and retrieval efficiency, and lower switching time and SD of entropy), and factor scores (red; higher factor 2 and lower factor 1 scores). **B-C.** The figures show the correlation between actual values and 10-fold cross validated support vector machine predictions of these values based on behavioral measures (green), BMAPS measures (blue), and factor scores (red). See Figure 3 for BMAPS parameter loadings on five factors. See **Table 1** for description of BMAPS variables.

Using cross-validated regression SVM (**Figure 6B-C**), we found that BMAPS-derived factor scores provided the best predictions of Numerical Operations (r = 0.34, p = 0.0012) and Mathematical Reasoning (r = 0.34, p = 0.0013), outperforming predictions based on the original model parameters (r 0.22 to 0.27) or behavioral measures alone (r 0.08 to 0.17). Stepwise regressions (**Tables S7-8**) revealed that baseline switching time uniquely predicted both Numerical Operations and Mathematical Reasoning, highlighting its potential importance as a cognitive marker of individual differences in mathematical achievement.

### BMAPS reveals distinct pathways linking affective factors to math performance

Next, we sought to evaluate the mechanisms by which math anxiety and positive attitudes influence problem solving. Math anxiety was assessed using the Scale for Early Mathematics Anxiety (SEMA)^46^. The SEMA included two subscales: numerical calculation anxiety and situational performance anxiety^46^. Positive math attitudes were assessed using the Positive Attitude Toward Math (PATM)^47^. SEMA calculation anxiety was negatively correlated with retrieval efficiency (r = -0.29, p = 0.035) and PATM was negatively correlated with baseline switching time (r = -0.30, p = 0.043).

We fit a series of structural equation models testing the direct and indirect (mediated by model parameters) effects of these affective factors on performance. **Figures 7A-C** show how the two SEMA subscales and PATM are directly associated with task-specific behavioral measures including accuracy on answered problems (AC), median reaction time (RT), and no-response rate (NR). To test for candidate mediating mechanisms, we evaluated multiple structural equation models that tested each of the BMAPS measures as potential mediators (**SI Table S9)**. A combined structural equation model (**Figure 7D**) revealed distinct pathways, with math anxiety impacting arithmetic task performance through retrieval efficiency, while positive math attitudes influenced arithmetic task performance through both retrieval efficiency and baseline switching time. After accounting for mediation paths, there were no residual significant direct effects of math anxiety and math attitude on the behavioral measures. This dual-pathway model showed good fit indices (*χ*^!^ p = 0.53 (should be > 0.05), CFI 1.00, TLI 1.008, RMSEA 0.000). An alternative control model specification (**Supplementary Figure S3**) with reversed links where affective factors (anxiety and attitude) were modeled as outcomes of the cognitive and behavioral measures failed to reveal any significant links between cognitive/behavioral measures and affective outcomes. These results provide new insights into the cognitive mechanisms by which affective and motivational factors contribute to problem-solving performance.

**Figure 7.**
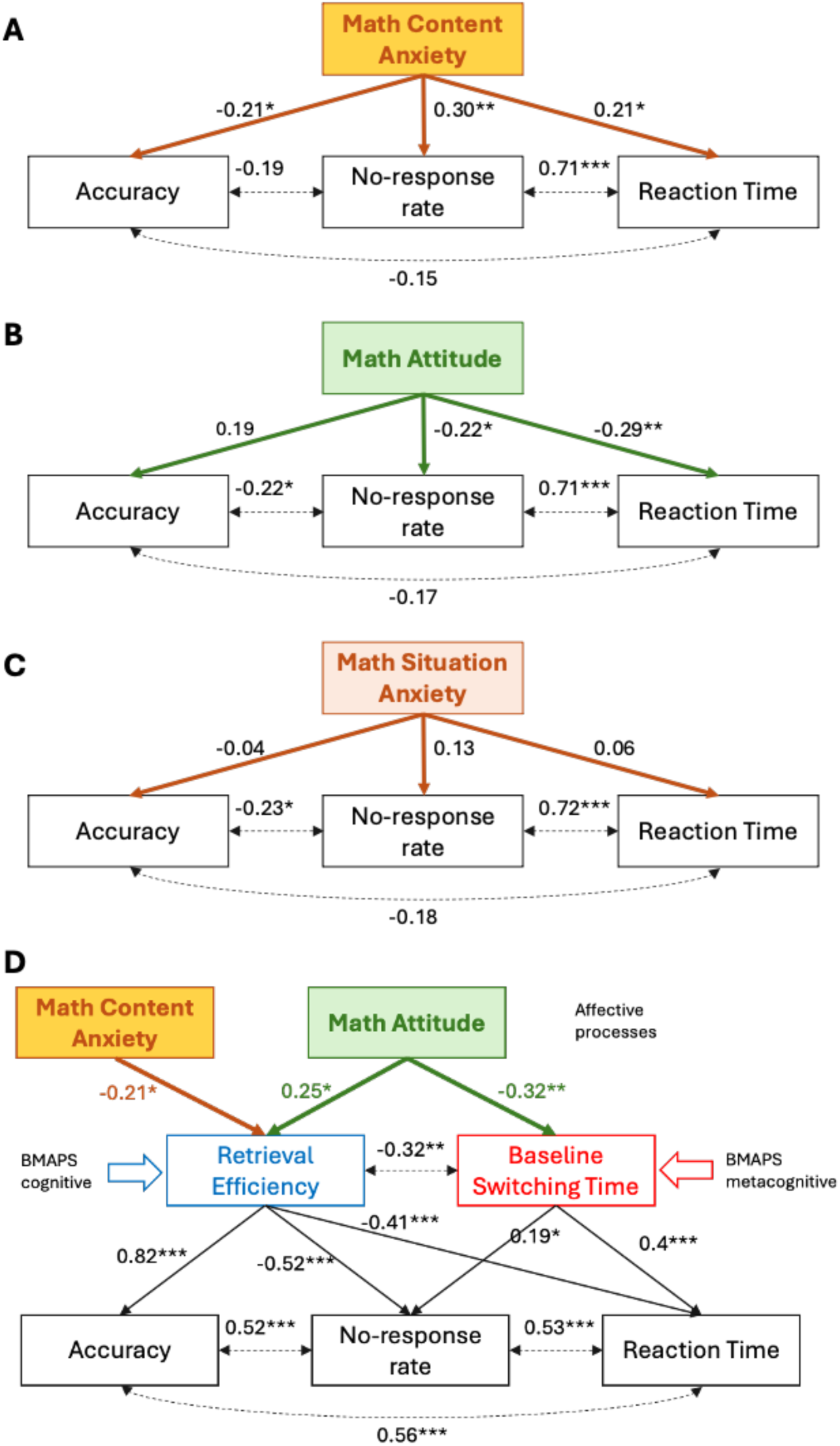
**A-C. Structural equation model (SEM) shows relationships between math attitude, math anxiety, and arithmetic task performance.** Math Attitude, Math Content Anxiety and Math Situation Anxiety (see *Methods* for description of variables) were related to behavioral measures of accuracy (AC), reaction time (RT), and no-response rate (NR) on the arithmetic task. Values on single sided arrows represent regression coefficients and those on double-sided arrows indicate covariances. **D. A multiple mediation model demonstrates pathways between math attitude, math anxiety, and arithmetic task performance explained by latent cognitive model parameters.** The relation between math anxiety and arithmetic task performance was mediated by retrieval efficiency, whereas the relation between math attitude and arithmetic task performance was mediated by both retrieval efficiency and baseline switching time. * p<0.05, **p<0.01, ***p < 0.001.

### Longitudinal BMAPS analysis identifies developmental shifts in cognitive processes

To examine developmental changes in problem-solving processes, we conducted a longitudinal analysis on a subsample of 30 participants assessed 1-2 years apart (mean ages 8.3 and 9.3 years). We observed significant changes in retrieval efficiency (t = 3.97, p = 0.0039), baseline switching time (t = -4.49, p = 0.0011), and decision threshold (t = -4.76, p = 0.0005), corrected for multiple comparisons (**Figure 8A-C, SI Table S10**). Specifically, retrieval efficiency increased, while decision threshold and baseline switching time decreased over the 1-2 years of development, consistent with expected gains in memory retrieval and cognitive control during this period. The mix of strategy use (retrieval versus procedural strategies) did not significantly change, based on both the cognitive model inferences as well as self-reports (*p*s > 0.05).

**Figure 8:**
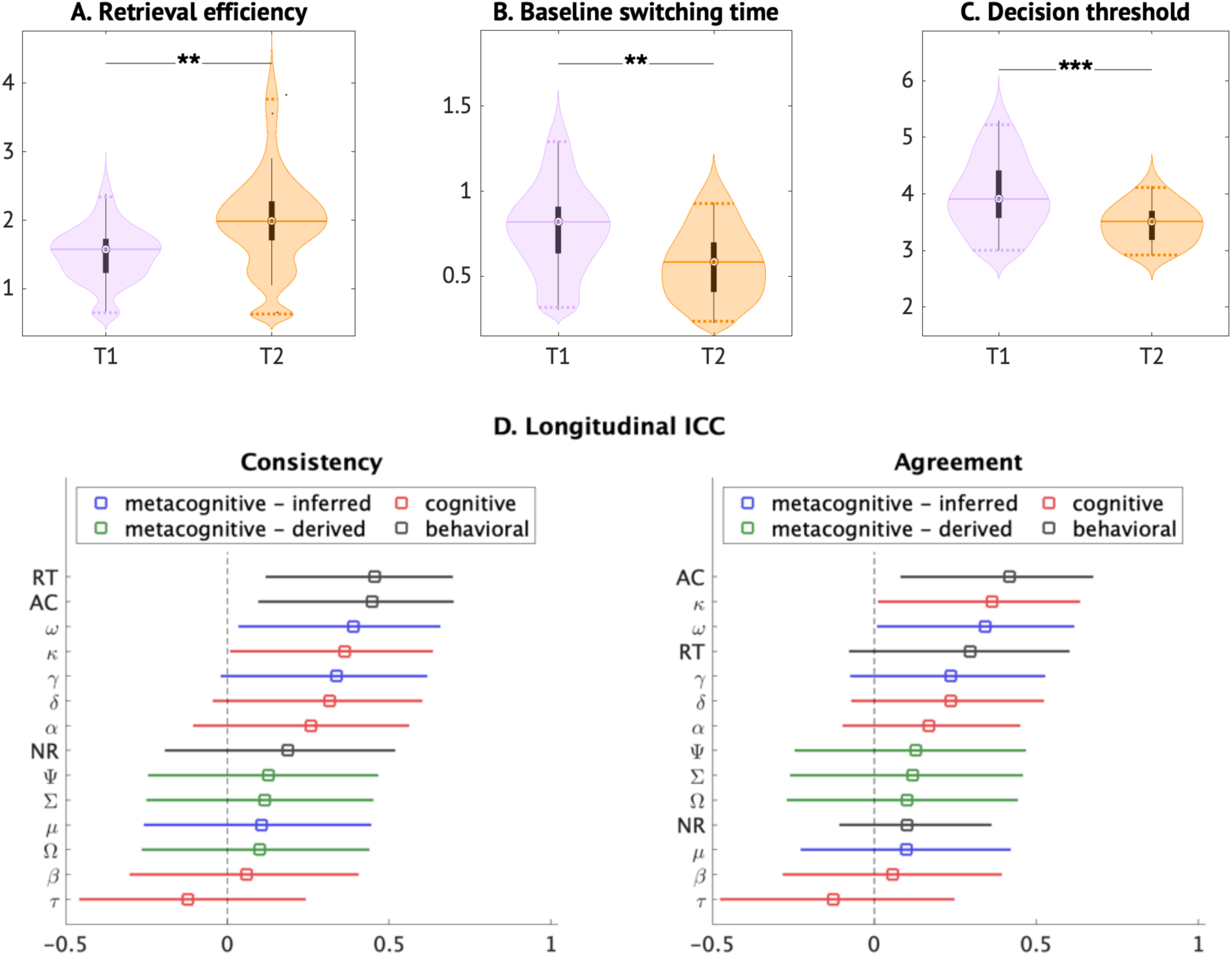
**Longitudinal analysis. A-C. Distributions of BMAPS parameters at first (T1) and second (T2) time points**. Significant longitudinal changes in group-level distributions were observed in three BMAPS parameters: retrieval efficiencies increased, while baseline switching times and decision thresholds decreased across two time points. These longitudinal changes indicate improved problem-solving capabilities across development. **D.** Longitudinal ICCs assessed levels of consistency and agreement between two time points for reaction time (RT), accuracy (AC), and no-response rate (NR), and BMAPS parameters. See **Table 1** for description of BMAPS variables. * p<0.05, **p<0.01, ***p < 0.001.

Baseline switching time remained a stable predictor of broad standardized Mathematical Reasoning WIAT-II-based performance across the two timepoints (**SI Tables S8, S11**). However, an analysis of within-subject changes indicated substantial variability in the rates of change across individuals for all model parameters and behavioral measures (**SI Table S12**), suggesting heterogeneous developmental trajectories (**Figure 8D**).

### BMAPS-based clustering reveals distinct problem-solving profiles

To determine whether our model could identify meaningful subgroups of problem solvers, we performed k-means clustering on participants’ BMAPS-derived factor scores. This analysis revealed three distinct clusters that differed systematically in their patterns of strategy use, problem-solving performance, and mathematical achievement (**Figure 9, Supplementary Table S13**).

**Figure 9:**
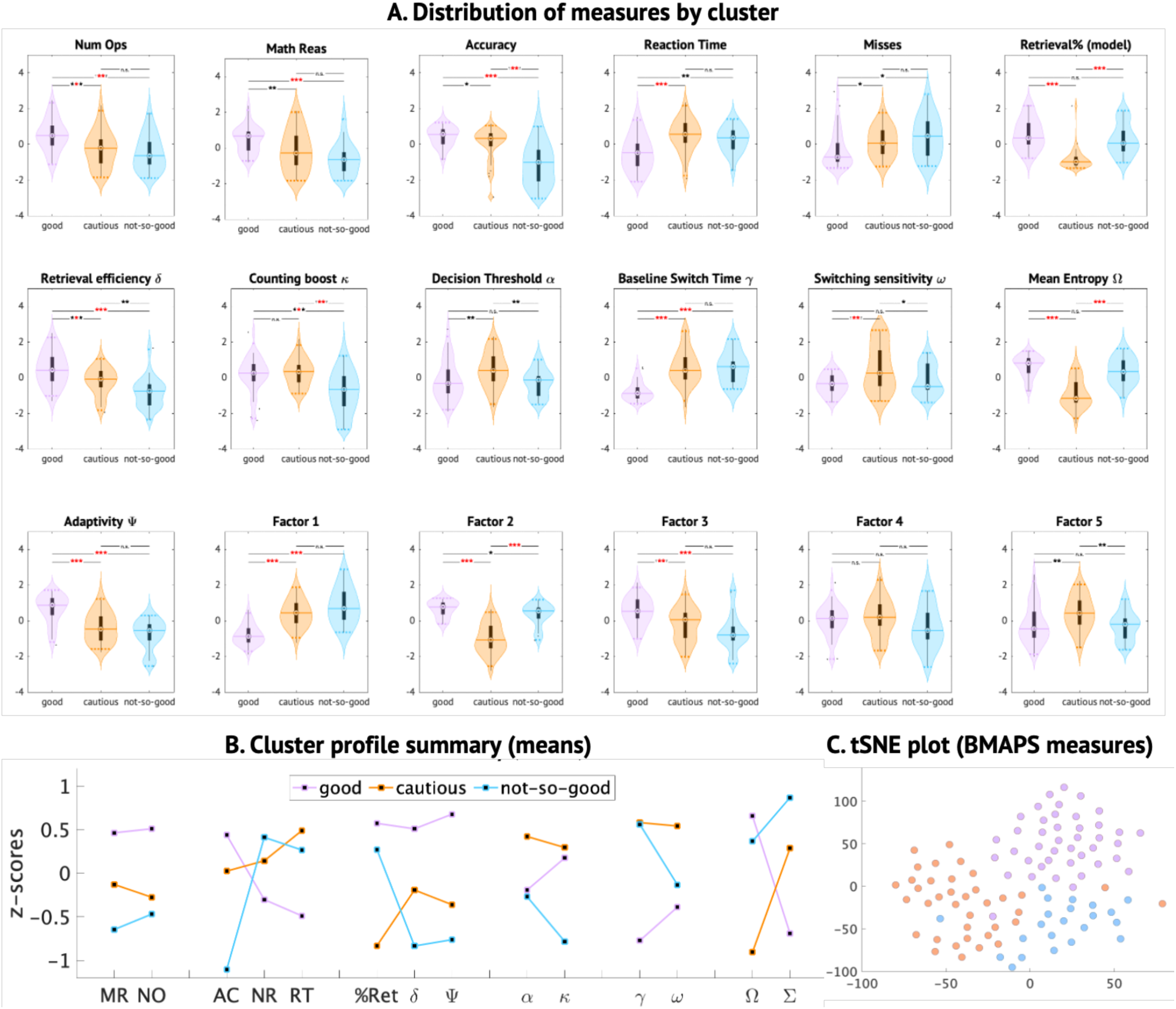
Clustering identified three distinct clusters associated with BMAPS parameters. The distribution of BMAPS parameters grouped children into “good-solvers” (pink), “perfectionist-solvers” (orange), and “not-so-good solvers” (blue) students. **A.** Distribution of measures of mathematical achievement, arithmetic task performance, BMAPS measures, and factor scores for the three clusters identified. **B.** Cluster means for key behavioral and BMAPS measures. **C.** t-Distributed Stochastic Neighbor Embedding (tSNE) plot showing three clusters. See Figure 3 for BMAPS parameter loadings on five factors. See **Table 1** for description of BMAPS variables. * p<0.05, **p<0.01, ***p < 0.001.

The “good solvers” cluster showed the highest retrieval strategy use, retrieval efficiency, and adaptivity, fastest baseline switching time, as well as the best performance on the arithmetic task and standardized WIAT-II subtests. The “not-so-good solvers” cluster had the lowest retrieval efficiency and adaptivity despite relatively high retrieval use, and performed poorly on the arithmetic task. The “perfectionist solvers” cluster had the lowest retrieval use, but the highest decision thresholds and counting strategy boost, and showed moderate performance across measures (**Figure 9A-C, Supplementary Figure S4**). These profiles align with and extend previous theoretical taxonomies of strategy development ^17,48^, demonstrating the utility of our model-based approach for identifying meaningful individual differences and sources of heterogeneity.

ANOVA of longitudinal changes in parameters within each cluster (**SI Table S14)** reveals that retrieval propensity (F = 6.55, p = 0.049) and mean entropy (F = 11.39, p = 0.003) showed a significant difference between groups. The “perfectionist solvers” show meaningful changes in the direction of “good solvers”. Among children who do not develop strong retrieval skills early during development, the children who are cautious about retrieval and have stronger dependencies on procedural or counting strategies (“perfectionist”) also show stronger retrieval strategy skills longitudinally, compared to children who use retrieval sub-optimally (“not-so-good”) earlier during development.

## Discussion

We developed a novel Bayesian model of arithmetic problem solving (BMAPS) to identify the latent cognitive and metacognitive processes underlying children’s mathematical abilities. Our model successfully captured dynamic variations in strategy use and efficiency across trials, providing a comprehensive view of the cognitive mechanisms that drive individual differences in performance and learning. The key findings are the following: First, multiple BMAPS cognitive and metacognitive parameters, including retrieval efficiency, strategy adaptivity, and switching time, collectively explained significant variance in problem-solving accuracy, response times, and error rates. Second, BMAPS parameters, especially retrieval efficiency and baseline strategy switching time, predicted individual differences in standardized measures of mathematical achievement above and beyond aggregate performance measures. Third, math anxiety and math attitudes influenced problem-solving performance through distinct pathways, with anxiety affecting retrieval efficiency and attitudes affecting strategy adaptivity. Fourth, longitudinal analyses revealed developmental improvements in retrieval efficiency and strategy adaptivity, with baseline strategy switching time emerging as a stable predictor of mathematical competence over time.

Our findings provide new insights into the complex interplay of cognitive and metacognitive skills and affective and motivational factors that shape children’s mathematical problem solving. By isolating specific mechanisms that contribute to individual differences and developmental changes, our model offers a powerful tool for advancing theory and informing educational practice.

### Uncovering latent metacognitive processes in mathematical problem solving

Our study takes a significant step forward in uncovering the latent metacognitive processes that underlie children’s problem-solving performance and strategy use. While previous research has highlighted the importance of metacognition in academic learning^49–51^, few studies have attempted to formally model and quantify metacognitive processes in the context of arithmetic problem solving ^18,25^. This is a critical gap, as metacognitive skills are thought to play a key role in the development of conceptual changes^52^ and adaptive expertise ^53,54^, allowing individuals to flexibly select and apply appropriate strategies in the face of changing task demands.

We define metacognition as the processes involved in monitoring and controlling one’s own cognitive processes during problem solving^55^. This includes the ability to accurately demonstrate knowledge or awareness of the effectiveness of different strategies for a given problem (monitoring) and to adaptively select and switch between strategies based on this assessment (control). Although these processes are not directly observable, BMAPS allows us to make inferences about latent metacognitive variables from observable behavioral data.

BMPAS operationalizes metacognition through several key model parameters and derived measures. First, retrieval propensity (μ) captures an individual’s overall tendency to select a retrieval strategy over other strategies, reflecting their metacognitive preference or bias. Second, baseline switching time (γ) represents the average time it takes an individual to switch between strategies, indexing their metacognitive efficiency in strategy selection. Third, switching sensitivity (ω) reflects how much an individual’s switching time is influenced by problem difficulty, capturing their metacognitive sensitivity to task demands. Fourth, adaptivity (ψ) quantifies the optimality of an individual’s retrieval strategy selection, based on the correlation between their retrieval efficiency and their actual strategy choices across trials. Higher adaptivity scores indicate better metacognitive calibration of strategy selection to one’s own abilities. Fifth, entropy (Ω and Σ) captures the degree of uncertainty or variability in an individual’s strategy selections across trials, with Ω reflecting the average entropy and Σ reflecting the variability in entropy. Higher entropy values indicate greater metacognitive flexibility or indecisiveness.

Crucially, latent metacognitive parameters showed high between-session reliability (ICC), thus representing potential computational phenotypes^56^ for individual differences in mathematical learning and problem solving.

Together, these metacognitive parameters allow us to quantify, for the first time, individual differences in dynamic strategy selection processes across time, which are hypothesized to play a critical role in adaptive problem solving. By incorporating these metacognitive components, our computational model enabled us to provide a comprehensive account of cognitive mechanisms underlying children’s arithmetic performance and learning.

### Cognitive and metacognitive underpinnings of mathematical problem-solving performance

Our first aim was to evaluate whether the cognitive and metacognitive parameters identified by BMAPS could explain individual differences in problem-solving performance. We found that retrieval efficiency, decision thresholds, strategy adaptivity, and baseline switching time were key predictors of accuracy, response times, and error rates. This suggests that the ability to efficiently retrieve arithmetic facts from memory and adaptively select strategies based on problem features are critical components of problem-solving success.

Importantly, our model-based approach allowed us to dissociate these cognitive processes with greater precision than previous approaches that rely on aggregate measures of performance. By estimating strategy use and efficiency at the trial level, our models captured fine-grained variations in performance. Moreover, by specifying distinct parameters for the execution and selection of strategies, the models identified unique contributions of these processes to individual differences in performance.

### Cognitive predictors of task-distal mathematical achievement

Our second aim was to assess the power of BMAPS parameters to predict individual differences in standardized measures of mathematical achievement. As hypothesized, we found that students with higher retrieval efficiency and strategy adaptivity showed advantages not only in arithmetic computation but also in broader assessments of mathematical concepts and problem solving using the WIAT-III Numerical Operations and Mathematical Reasoning subtests. Control analysis showed that BMAPS measures and factor scores were predictive of domain-specific broad mathematical achievement scores, but not of general cognitive abilities indexed by IQ. This suggests that the cognitive skills captured by our model are foundational for the development of mathematical proficiency.

Notably, the model parameters outperformed aggregate performance measures in predicting achievement scores. This demonstrates the added value of a process-level approach for understanding the cognitive bases of mathematical competence. By identifying specific mechanisms that support learning and transfer, our model can help researchers and educators develop more effective methods for assessing and promoting mathematical skills.

These findings also have implications for theories of mathematical development. They suggest that the acquisition of efficient retrieval and adaptive strategy selection may be key milestones in the progression from novice to expert problem solving. By tracking these cognitive processes over time, researchers can gain new insights into the trajectories of skill acquisition.

### Affective and motivational influences on problem-solving competency

Our third aim was to leverage BMAPS to elucidate the specific pathways by which math anxiety and math attitudes influence problem-solving performance. Such measures have been hypothesized to be strong predictors of academic achievement^57^. However, it is not known how math anxiety and positive math attitude relate to math achievement, and whether they affect performance through common or dissociated functional paths. Using structural equation modeling, we found that math anxiety impacted performance primarily via cognitive processes by reducing retrieval efficiency, while positive attitudes promoted performance both, via cognitive (improving retrieval efficiency), and metacognitive (reducing baseline switching time), processes (**Figure 7**). These results provide a more precise understanding of the mechanisms by which affective and motivational factors shape cognitive processing and problem solving.

Previous research has documented the negative effects of math anxiety on strategy use and arithmetic fluency^42,43^, but the underlying mechanisms have been unclear. Our findings suggest that anxiety may disrupt the retrieval of arithmetic facts from memory, leading to slower and more error-prone performance. In contrast, we found that positive attitudes towards math promoted performance by increasing the adaptivity of strategy selection. Students with more positive attitudes were better able to calibrate their strategy choices to problem features, suggesting greater metacognitive awareness and control. This aligns with research showing that positive academic emotions can facilitate flexible problem solving^47^.

Importantly, our model allowed us to test the directionality of these effects by comparing alternative path specifications. The best-fitting model indicated that anxiety and attitudes influenced cognitive processing, rather than the reverse. This supports the idea that affective factors can shape the development of cognitive skills, consistent with theories of emotion-cognition interactions in learning^58,59^.

These findings have practical implications for education, as they suggest potential targets for interventions aimed at improving mathematical performance. Strategies that reduce math anxiety, such as cognitive reappraisal, may help students develop more efficient retrieval skills. Similarly, interventions that foster positive attitudes towards math, such as growth mindset training^60,61^, may promote the development of adaptive strategy selection. By addressing both cognitive and affective factors, educators can create more supportive learning environments that enable all students to reach their full potential.

### Developmental dynamics of strategic expertise

Our fourth aim was to examine how problem-solving strategies change over the course of learning and development. Using longitudinal data, we mapped the trajectories of strategy use and efficiency across one-two years of schooling. We found that children showed significant improvements in retrieval efficiency and strategy adaptivity over time, with larger gains associated with greater improvements in problem-solving accuracy and mathematical achievement.

Notably, we identified baseline strategy switching time γ as a key predictor of growth in mathematical competence. Students who were faster at switching between strategies at the first timepoint showed greater gains in retrieval efficiency, strategy adaptivity, and overall performance over time. This suggests that the ability to flexibly shift between strategies may be a foundational skill that supports the development of strategic expertise^56^. Findings also reveal the importance of considering developmental clusters and heterogeneous growth trajectories for different aspects of cognitive and metacognitive processes over development.

These findings extend previous research on the development of problem-solving strategies ^11,17,18,22,62,63^ by providing a more detailed view of the cognitive processes that underlie growth. By tracking changes in specific model parameters over time, we can identify the mechanisms that drive improvements in performance and the factors that constrain or enable progress. This approach can inform the design of educational interventions that are tailored to the developmental needs of individual learners.

The longitudinal results also have implications for theories of cognitive development. They suggest that the acquisition of strategic expertise involves the coordination of multiple cognitive processes that develop at different rates over time. By examining the interplay of these processes and their relations to academic outcomes, we can gain new insights into the factors that support or hinder learning.

### Quantifying a theoretical model of variability in strategy use and efficiency

Our findings provide a quantitative and mechanistic understanding of variability in strategy use and efficiency, extending Siegler and Shipley’s theoretical model of strategy choice^64^. They identified key characteristics of adaptive variability in children’s strategy selection during arithmetic problem solving, including dynamic variability within individuals, context-sensitive adaptivity, changes in strategy use and efficiency over time, generalizability of acquired strategies, and individual differences reflecting distinct cognitive styles^16,64,65^. However, previous studies relied on verbal self-reports to assess strategy use, which can be less accurate or feasible, particularly in children^30–34^, and no comprehensive quantitative models have been developed to capture these complex dynamics.

Our study rigorously quantifies and extends several aspects of strategy choice models for the first time without relying on self-reports from participants. In BMAPS, the decision threshold parameter α captures the confidence criterion, while baseline switching time γ and switching sensitivity ω denote the search length prior to strategy shifts. Entropy Ω quantifies variability of strategy mixtures within individuals, adaptivity index ψ indexes optimal strategy selections, and retrieval efficiency δ represents retrieval execution efficiency. Thus, BMAPS provides a rigorous quantification of the key components of Siegler and Shipley models which proposed that a confidence criterion must be surpassed for retrieval strategy use, along with a search length parameter denoting the duration of retrieval attempts before shifting to a backup strategy^16,64,65^.

Crucially, this quantification allowed us to investigate individual differences and heterogeneity in strategy use and efficiency. Clustering observable performance measures (**Figure 9**) identified three subgroups: (i) good students, frequently using fast and accurate retrieval; (ii) cautious low- retrievers, efficiently using retrieval but underusing it compared to good students; and (iii) not-so-good students, inefficiently using retrieval. Good students used retrieval 6% more than not-so-good students. These subgroups resemble those proposed by behavioral strategy choice models^17^ ^48^, providing concurrent validation of decades-old qualitative models while offering unsupervised, multidimensional, quantified individual differences measures without relying on self-reports. This showcases the power of our computational approach for uncovering meaningful patterns of variability in children’s strategic problem solving.

### Integrative insights from the BMAPS cognitive model

In addition to testing our main hypotheses, we conducted a series of analyses to validate the cognitive model and explore its broader implications. These analyses yielded several integrative insights that enhance our understanding of the complex dynamics of mathematical problem solving.

First, the model’s ability to recover meaningful individual differences and strategy-related performance patterns demonstrates its construct validity. By clustering students based on their model parameters, we identified distinct problem-solving profiles that aligned with previous theoretical taxonomies^16,64,65^ ^17^. This suggests that our model captures key dimensions of variation in children’s strategic behavior and provides a useful framework for characterizing individual differences.

Second, the model’s predictive power and reliability highlight its potential for educational applications. The fact that model parameters outperformed aggregate performance measures in predicting achievement scores suggests that process-level assessments may be more informative for identifying students at risk for difficulties. Moreover, the higher stability of metacognitive model parameters across sessions indicates that they capture meaningful traits that can be targeted for intervention. By identifying the specific cognitive skills that underlie problem-solving success, our model can inform the design of targeted interventions and support systems. For example, students who struggle with fact retrieval may benefit from different instructional approaches than those who have difficulty adapting their strategy use to problem types. By tailoring interventions to individual profiles of strengths and weaknesses, educators can more effectively support the development of mathematical competence. Moreover, BMAPS ability to quantify theoretical constructs such as strategy adaptivity and switching time provides a bridge between cognitive theory and educational practice. By operationalizing these constructs in terms of specific parameters, our model can help researchers test and refine theories of strategic development. In turn, this can inform the design of instructional methods and assessment tools that are grounded in cognitive science.

## Conclusions

Our study presents a novel computational approach for understanding the cognitive and metacognitive processes that underlie children’s mathematical problem solving. By developing a Bayesian model that captures trial-by-trial variations in strategy use and efficiency, we were able to identify specific mechanisms that drive individual differences in performance and learning.

Our findings highlight the importance of retrieval efficiency and strategy adaptivity as foundational skills for mathematical competence, and suggest promising avenues for educational interventions that target these capacities.

More broadly, our study demonstrates the power of integrating cognitive modeling with educational research to advance both theory and practice. By providing a precise and comprehensive view of the cognitive processes that support learning, models such as those developed in our study can inform the design of assessments and interventions that are tailored to the needs of individual students. In turn, educational settings provide rich opportunities to test and refine cognitive theories, leading to a virtuous cycle of scientific discovery and practical application.

## Methods

The combination of computational modeling, longitudinal analysis, and multivariate statistical techniques allows for a comprehensive investigation of cognitive and metacognitive mechanisms underlying children’s arithmetic problem solving and their relation to academic achievement, affective and motivational factors, and developmental change. **Figure 1** provides an overview of our study methods and analysis pipeline.

### Participants

132 children in 2nd and 3rd grades (ages 7-9) recruited from Bay Area schools were recruited as a part of a larger longitudinal study. All participants had the Wechsler Abbreviated Scale of Intelligence (WASI^66^) Full-scale IQ scores greater than 80, and had no history of major psychiatric or neurological disorders. Among them, 105 (46 females; mean age=8.73 years). participants were included in the final analysis as they had complete data for three separate sessions and met the threshold for having at least 50% accuracy on binary choice problems in each of these three sessions. Informed written consent was obtained from the legal guardian of the child and all study protocols were approved by the Stanford University Institutional Review Board. All participants were treated in accordance with the American Psychological Association Ethical Principles of Psychologists and Code of Conduct.

### Arithmetic problem-solving task

Children solved addition problems (e.g., 3+4=7) in a verification format during fMRI scanning. Participants were asked to respond with button press 1 for correct answers and button press 2 for incorrect ones. Problems varied in size, with sums ranging from 3 to 14. Details of the task are described elsewhere^67^ and in the **Supplementary Information**.

### Strategy assessment task

In a separate session, children reported strategies used to solve addition problems. Strategies were classified as retrieval, counting, or decomposition. This data was used to validate BMAPS- inferred strategies. Details of the task and methods to determine and validate these strategies are described elsewhere^27^ and in the **Supplementary Information**.

### Standardized cognitive and academic achievement measures

Children completed Wechsler Individual Achievement Test, Second Edition (WIAT-II ^68^) Numerical Operations and Math Problem Solving subtests, assessing arithmetic fluency and math reasoning. These measures were used to assess broad mathematical reasoning. IQ was assessed using the WASI^66^.

### Math anxiety and attitude to math

Math anxiety was assessed using the Scale for Early Mathematics Anxiety (SEMA), which includes two subscales: numerical calculation anxiety and situational performance anxiety^46^. Math attitudes were measured using Positive Attitude Toward Math (PATM) and calculated as positive attitudes towards math relative to other subjects^47^. Further details are included in the **Supplementary Information.**

### BMAPS model

We developed a novel Bayesian model that models cognitive and metacognitive processes and infers the probability of using retrieval, counting, and decomposition strategies on each trial based on response times and accuracy (**Figure 2**). BMAPS also estimates individual-level parameters related to strategy execution (retrieval efficiency, counting ability) and strategy selection (retrieval propensity, switching time, adaptivity, entropy). Details on the mathematical specification of the Bayesian model are provided in the **Supplementary Information**.

### Factor Analysis

Factor analysis was carried out in MATLAB using Promax rotation. To compare and evaluate the adequacy of factor analysis models, we used the following criteria: CFI (comparative fit index; threshold 0.95), TLI (Tucker-Lewis index; threshold 0.95), and RMSEA (threshold < 0.08).

### Individual differences analysis

We used canonical correlation analysis (CCA) to identify multivariate relationships between model parameters and arithmetic performance measures. We also used regularized regression and SVMs to predict WIAT-II scores from model parameters. Hierarchical clustering was applied to model parameters to identify subgroups of children with distinct problem-solving profiles. Other prediction modeling and related analysis is detailed in the **Supplementary Information**.

### Mediation analysis of affective factors

We used structural equation modeling to test hypothesized mediation effects of math anxiety and attitudes on arithmetic performance through model parameters. We compared alternative path configurations to assess the directionality of effects.

### Longitudinal analysis

A subset of children (N = 30) were re-tested on the arithmetic task and WIAT-II subtests 1-2 years later. We fit the BMAPS model to the first timepoint (T1), fixed the item-level parameters based on inferences from T1, and then fit BMAPS with the fixed item-level parameters to T2 data. Thus, longitudinal changes could be measured in individual level parameters while controlling for item effects across the two timepoints.

### Statistical analysis

All p-values were corrected for multiple comparisons, with original and adjusted p-values provided in the Supplementary tables.

## Acknowledgement

This research was supported by grants from the National Institutes of Health (HD094623, HD059205, MH084164) and National Science Foundation (DRL-2024856); and Stanford Maternal & Child Health Research Institute Postdoctoral Support Awards to PM and HC. We thank all participating children and their families.

