## Supplemental Information for "Unraveling latent cognitive, metacognitive, strategic, and affective processes underlying children’s problem-solving using Bayesian cognitive modeling"

### Supplementary Information

**I. Supplementary Methods**

**II. Supplementary Results**

**III. Supplementary Figures**

**IV. Supplementary Tables**

#### I. Supplementary Methods

***Arithmetic problem-solving task***

Children's arithmetic problem-solving skills were examined during 3 fMRI sessions (1 block design, 2 event design sessions). The block-design session included 36 problems, of which 18 were simple control problems (addition by 1) and 18 were complex addition problems, The 2 event-design sessions included 52 problems each, of which 26 in each were simple control problems (addition by 1) and 26 were complex addition problems. Each individual thus solved 70 complex addition problems and 70 simple control problems (addition by 1). In addition, the block-design also included symbol-finding and passive fixation trials. In the complex addition problems, participants were presented with an equation involving two addends and were asked to indicate, via a button press, whether the presented answer was correct (e.g., "3 + 5 = 8"). The first operand ranged from 2 to 9, the second operand ranged from 2 to 5, and tie problems were excluded. Correct answers appeared in 50 % of the trials. Incorrect answers deviated by ± 1 or ±2 from the correct sum. The simple addition problems were identical except that one of the addends was always '1' (e.g., "3 + 1 = 4"). Each stimulus was displayed for 5 seconds.

***Strategy assessment task***

In an independent strategy assessment task, each participant's strategies for solving addition problems were assessed using a standardized trial-by-trial assessment of strategies[1]. Participants were instructed to use whatever strategy was easiest for them to get the answer. Each problem was presented on the center of the computer screen. Participants were asked to solve each problem and state the answer out loud as quickly as possible. Participants were then asked to report how they solved each problem. For each problem, the experimenter took detailed notes of overt signs of counting, such as finger counting. Based on each participant's self-reported strategy and experimenter's observations, strategies for solving addition problems were classified into several categories of interest: 'counting' (e.g., counting fingers, verbal counting, counting in mind), 'retrieval' (e.g., remembering the answer to the problem), and 'decomposition' (e.g., break-apart). A set of 18 addition problems with two addends from 2 to 9 and sums ranging from 6 to 17 was used. Problems with two identical addends or with addends of 0 and 1 were excluded due to low strategy variability. Half of the problems were randomly presented in larger addend plus smaller addend format (e.g., "5 + 3") and the other half were presented in the reverse order (e.g., "3 + 5").

***Math Anxiety and Attitude to Math***

Math anxiety was assessed using the 20-item Scale for Early Mathematics Anxiety [2] (SEMA), which included two subscales: numerical calculation (content) anxiety (first 10 items) and situational performance anxiety (last 10 items). Numerical calculation anxiety scales assessed children's anxiety related to solving problems from second and third grade mathematics curriculum. Situational performance anxiety scales assessed children's anxiety related to social and testing situations that second and third graders may encounter while learning math concepts. For each item, children were asked to rate how anxious they felt on a five-point response. Children responded by selecting one of the faces graded from not anxious to anxious or by verbally replying how anxious or non-anxious they felt. An individual's total math anxiety (or numerical calculation anxiety or situational performance anxiety) score was computed by summing the 20 items' (or 10 items') ratings.

Math attitudes were measured using the 12-item Positive Attitude Toward Math (PATM) and calculated as positive attitudes towards math relative to other subjects[3]. Six questions measured two aspects of math attitudes, namely, strong interests (e.g., "How much do you like math?") and self-perceived ability (e.g., "How good are you at learning math?"), and six questions measured attitude toward general academic subjects. Participants were asked to respond on a 5-point Likert-type scale. A mean score averaging responses on the six math-related questions was divided by a mean score on the six general questions to estimate positive attitudes towards math relative to other subjects.

***Mathematical specification of the BMAPS model***

We developed novel process models of mathematical problem solving. To address the lack of methodological and computational rigor in previous work, we developed computational models that infer differences in the latent cognitive strategies used by an individual on a trial-by-trial basis, dissociate between multiple cognitive sub-processes, and account for sequential as well as item level effects. Our approach allowed us to investigate multi-dimensional representations of individual differences that provide a greater functional resolution of neurocognitive processes.

Basic decision process

The arithmetic tasks are primarily presented as an answer verification problem, with choice reflecting a binary yes/no response. The responses (choice – accurate or inaccurate, and reaction time) are modeled as a drift diffusion model (DDM). The response of an individual $i$ on trial $t$ is denoted as $x_{it}$ where $x_{it} = 1$ if the response was correct and $x_{it} = -1$ if the response was incorrect. The response time is similarly denoted as $r_{it}$. The choice coded response time is represented as $y_{it} = x_{it}r_{it}$. Here, $y_{it}$ is treated as a sample from random variable $Y_{it}$ which is distributed according to a Wiener distribution [4,5]. The Wiener distribution describes the probability density function for $Y_{it}$, or in other words, the joint pdf for the random variables $X_{it}$ and $R_{it}$. The Wiener distribution is a four-parameter distribution that describes the first passage time of the diffusion process, that is, the distribution of times taken for the diffusion process to first hit the positive ($x = 1$) and negative ($x = -1$) decision boundaries. This is a canonical approach for modeling the joint distribution of choice and response time in two-choice processes[6,7].

$$y_{i,t} \sim \text{Wiener}\left(\alpha_{i,t}, \tau_{i,t}, \beta_{i,t}, \delta_{i,t}\right) \quad (1)$$

Accounting for multiple strategies

Children as well as adults are however known to use multiple cognitive strategies (e.g. memory retrieval, counting, decomposition, etc.) to solve mental arithmetic problems. Each of these latent cognitive strategies can be represented as a unique Wiener process, with its own set of parameters characterizing the nature of the process. Behavior is thus modeled as a *mixture model* of these latent strategies across trials. A latent value codes which strategy produces a response on any single trial: *S(t)*. The latent mixture drift diffusion model implemented can then be represented as:

$$y_{i,t} \sim \text{Wiener}\left(\alpha_{i,S(t)}, \tau_{i,S(t)}, \beta_{i,S(t)}, \delta_{i,S(t)}\right) \quad (2)$$

A key novel approach in this paper is to define the structure of some of these parameters based on what we know about each strategy, and how strategy execution may depend on the characteristics of the problem (item type).

Process Dissociation: Strategy Selection

The above characterization defines the structure for some key arithmetic strategies, and defines the dependence of strategy execution of item characteristics, but does not specify how strategy selection takes place on each trial, for each item.

*A two-step strategy selection process:* There is some evidence for autonomous retrieval and serial processing of retrieval versus other computational strategies [8,9]. Our model assumes a two-step sequential process, with a primary response demanded via a memory retrieval strategy, and a possible subsequent switch to alternate strategies, such as counting or decomposition. The secondary choice between alternate strategies is considered in parallel. Similar two-step sequential response processing that involves an initial recognition step, possibly followed by a polytomous choice between multiple options (depending on the outcome of the recognition step) has been characterized by sequential IRT or SRM-MC models [10]. Whilst such models are typically used to characterize the response process when selecting between observable multiple choices, we adapt this class of models to characterize the process of sequential selection between latent strategies. Following this approach, the selection of strategies is assumed to be dependent on a sequential process, with the first step modeling whether there is persistence of memory retrieval. This probability is modeled as a traditional 1PL-IRT model. Thus, for step 1, the probability of an individual *i* selecting a memory retrieval strategy for an item $k$ ($s_{ik} = M$), depends on the individuals' proclivity towards memory retrieval $\mu_{iM}$ as well as how amenable the item is to be retrieved from memory $\eta_{kM}$ (this is measured at a group level), and follows the form of 1PL-IRT model. The $\eta_{kM}$ parameter can be interpreted as the ease of retrieval of item *k.* Importantly, this breaks down the propensity of retrieval into orthogonal individual and item level effects.

$$p(s_{ik} = M) = \frac{1}{1 + e^{-(\mu_{iM} + \eta_{kM})}}$$

In the second step, assuming that the result of the first step is to switch away from retrieval ($s_{ik} \neq M$), a nominal response model [11] is considered. This step models the probability of selecting an alternate strategy $S$ by simultaneously taking into consideration the remaining 2 alternate strategies (counting and decomposition), where the probability of counting depends on item level effects ($\eta_{kC}$):

$$p(s_{ik} = C \mid s_{ik} \neq M) = \frac{1}{1 + e^{-(\eta_{kC})}}$$

This sequential process implies some switching point at which an individual might give up on the retrieval strategy and switch to an alternate strategy, in other words, the point at which the second step is initiated. Our proposal characterizes this switching time ($\xi_{ik}$) as the point at which a decision is taken on whether or not to switch away from memory by individual *i* for item *k*. It is possible that the actual retrieval takes longer, but enough confidence has been gained by then to take a decision on whether or not to switch. This switching time will vary by individual, but it may further vary depending on the nature of the item. We propose that people may have some form of access to the ease of retrieval of an item $\eta_{kM}$ and their switching time may thus vary depending on a baseline switching time $\gamma_i$ , switching sensitivity $\omega_i$ , and ease of item retrieval $\eta_{kM}$.

$$\xi_{ik} = \gamma_i + \omega_i \, \eta_{kM}$$

<u>Defining the parameters each process – Accounting for item level differences</u>

*Memory retrieval process*

The memory retrieval process is characterized as a drift diffusion process, which posits that evidence accumulates over time resulting in a decision when a decision threshold is reached. The efficiency of evidence accumulation is characterized by a memory retrieval drift rate parameter ($\delta^M$), with higher values characterizing faster and more accurate retrieval. The decision threshold parameter (α) captures the degree of confidence required to conclusively decide that a solution is retrieved from memory, with higher values characterizing slower but more accurate retrievals. The memory retrieval drift rate for an individual *i* for an item *k* is proposed as a combination of individual ($\delta_i^M$) and item level effects:

$$\delta_{ik}^M = \frac{\delta_i^M}{1 + e^{-\eta_{kM}}}$$

Thus, for items which are considered very easy for retrieval (high positive values of $\eta_{kM}$), $\delta_{ik}^M \rightarrow \delta_i^M$, whereas for items considered very difficult for retrieval (high negative values of $\eta_{kM}$), $\delta_{ik}^M \rightarrow 0$.

Finally, in a verification task, where one answer is provided and the subject has to respond true/false, a bias parameter ($\beta_i^M$) can measure any bias towards or away from verification, taking on values from 0 to 1, with a value of 0.5 representing no bias, higher than 0.5 representing bias towards verifying a presented answer as correct and a value less than 0.5 representing bias towards verifying a presented answer as incorrect.

*Counting process*
The counting process is also characterized as a drift diffusion process, with a combination of individual and item level effects. The counting strategy used in this model is assumed to be *min* counting (counting up from the larger addend). For an item $k$, with the first addend presented being $n_{1k}$ and the second being $n_{2k}$, the drift rate for counting is modeled as follows:

$$\delta_{ik}^{C\,[min]} = \frac{\delta_i^C}{\min\,(n_{1k}, n_{2k})}$$

*Decomposition process*
The decomposition process can be characterized as a set of $n_D$ steps taken to break down the addition process. For instance, 8+7 may be broken down as 7 = 2+5; 8+2 = 10; 10+5 = 15, which involves $n_D = 3$ steps.

$$\delta_{ik}^D = \frac{\delta_i^D}{n_D}$$

Each of these steps is at least as difficult as a counting step, so we apply the constraint, $\delta_i^D \leq \delta_i^C$

Derived Parameters
For adaptivity, we estimated the correlation across trials between latent memory retrieval efficiency and probability of selecting memory retrieval use on each trial. Higher adaptivity indicated greater correspondence between retrieval efficiency and use of retrieval strategy. For entropy, we measured the latent probability of selecting each of the three strategies (retrieval, counting, decomposition) on each trial. Higher entropy indicated that the individual was more uncertain about their strategy choice. The SD of entropy indicated the variability in the degree of uncertainty in strategy choice across trials.

***Model implementation and parameter recovery***
The models were implemented in a hierarchical Bayesian framework in JAGS [12] which implements a Gibbs sampler for Markov Chain Monte Carlo (MCMC) simulations. The following MCMC sampling hyperparameters were used: Total 3 chains with 70000 samples each. For each chain, we had a burn-in of 45000 samples, and a thinning factor of 5 was used to record 5000 of the remaining 25000 samples, based on which final inferences were made. Convergence was assessed using R-hat <= 1.1 across all chains. We also conducted a parameter recovery analysis by simulating behavior with the BMAPS model ad testing recovery of the generating parameters. Parameter recovery is not an assessment of the validity of the BMAPS model or its assumptions, nor a measure of effectiveness of the Bayesian methods used to make inferences. It does provide a way to check the implementation of the model, diagnose any potential identifiability issues, and understand the adequacy of the experimental designs for making useful model inferences. The model showed strong parameter recovery, with median correlation between generating and recovered parameters r = 0.87 (correlations ranged from 0.53 to 0.97, with all but 1 parameter having correlations > 0.75).

***Retrieval prediction***
Unsupervised BMAPS model based predictions: The average retrieval use inferred by the cognitive model based on in-scanner behavioral data is taken as prediction for the out-of-scanner retrieval data. This is completely unsupervised since the model does not ever have access to the out-of-scanner data.

Supervised predictions: We use 10-fold cross-validation, a regular least squares regression using a SpaRSA solver and lasso regularization and test two models to predict self-reported retrieval use. The first used parameters from the unsupervised BMAPS model (the supervised aspect refers to training the regression model) as predictors. The second model used behavioral performance (accuracy, RT, no response rates) and neuropsychological measures as predictors.

RT model (control model for comparison) predictions: This model assumes that reaction-time is a reliable discriminator of strategy use, which is a procedure commonly used for strategy classification in studies of arithmetic cognition. Using a 10-fold cross-validation procedure, we select the optimal cut-off reaction-time for each of the 10-fold subsets that minimizes the root mean square error between training data and predictions, where the predictions are made such that all trials with reaction times lower than the cut-off are classified as retrieval, and all others as non-retrieval.

***Permutation tests for assessing the range of within-individual variability in self-reports***

Since the out-of-scanner strategy assessment is only a proxy ground truth for the in-scanner retrieval data, we would expect a natural level of variability in how well these two situations match up. We ran a permutation analysis by selecting subsamples of the out-of-scanner strategy assessment, varying the analysis from 4-fold to 12-fold (the full set had 24 problems), and in each cross-validation, the actual out-of-scanner average retrieval from the training set was used to predict the retrieval of the test sets. The performance metrics from these permutations were captured as empirical within-individual permutations, and provide a tolerance benchmark for other methods. In other words, we expect the performance metrics from good methods of prediction to fall somewhere within the range (or better) provided by the within-individual permutation, since predicting from in-scanner to out-of-scanner itself is a form of within-individual permutation. As a contrast, a between individual permutation is performed by scrambling individuals and using out-of-scanner strategy reports from random individuals to predict other individuals. This provides a low end of the benchmark signifying that performance metrics falling within this range of between-individual permutation signal prediction methods that are as good as random guessing.

#### II. Supplementary Results:

***SI: Meaningful differences exist between within-participant variability and between-participant variability in use of retrieval versus procedural strategies – providing a reference point for evaluating face validity of independent inferences***

Children use a mix of different strategies depending on the problem, and often use a mix of different strategies even when solving the same problem multiple times. It is hence difficult to precisely validate inferences about which strategies were used on a particular trial against strategies used on a different set of problems or in a different session. This is further compounded by the fact that self-reports of strategy use may not be perfectly accurate for individual problems. However, if self-reported within-individual variability of strategy use across problems is reliably lower than between-individual variability, this provides us with reference levels to evaluate the face validity of independently inferred strategy use. We looked at cross-validated predictions about average strategy use by individuals on a subset of problems based on a different set of problems (within-participant predictions), and predictions about strategy use based on permuting individuals (between-participant predictions). As expected, within-participant predictions were significantly stronger than between-participant predictions as measured by unadjusted Kendall's tau (effect size 4.288; mean tau 0.283 95% CI from 0.071 to 0.522 for within, versus mean tau -0.229, 95% CI from -0.384 to 0.050 for between), and mean squared error (effect size -5.471; mean MSE 0.062, 95% CI from 0.028 to 0.099 for within versus mean MSE 0.194, 95% CI from 0.147 to 0.242 for between). These 95% CI ranges, and the effect size provide a strong reference for evaluating other independent inferences about strategy use – those that are similar to within-participant and different from between-participant predictions would demonstrate strong face validity.

***SI: Unsupervised cognitive model inferences of individual level in-scanner retrieval strategy use show face validity***

We compared unsupervised estimates about each individual's retrieval strategy use in-scanner as inferred by the cognitive model, with the self-reported use of retrieval in independent out-of-scanner sessions. The in-scanner sessions had 22 problems and the out-of-scanner sessions had 24 problems, with 11 problems common between the two sessions. Treating the unsupervised inferences across all problems as predictive of the average out-of-scanner strategy use, we obtained Kendall's tau 0.1934 and MSE of 0.0944, both within the 95%CI for within-participant variability but outside the 95%CI for between-participant variability.

***SI: Cognitive model parameters provide better predictions of retrieval strategy use compared to predictions based on behavioral and neuropsychological measures***

We used the parameters from the unsupervised cognitive model (which made unsupervised inferences about in-scanner problems) in a linear regression model to predict the strategy use for out-of-scanner problems. We compared this to a linear regression model using children's behavioral measures (accuracy levels, median RTs, proportion of missed problems) and neuropsychological measures (numops, math reasoning, and 4 WMTB-C measures). The model based on cognitive model parameters had better fit (tau 0.3260 vs 0.2896 and MSE 0.0533 vs 0.0585), and was more robust after cross-validation (mean tau 0.231 vs 0.146, effect size 1.923, $p<0.0001$and mean MSE 0.067 vs 0.076, effect size -1.963, $p<0.0001$). Comparing the cross-validation iterations for each of the models against the range of within-participant and between-participant measures, the absolute effect size for difference with the within-participant predictions was smaller for the cognitive model (Cohen's d for tau 0.493 versus 1.286, and for MSE -0.321 vs -0.917). Similarly, the absolute effect size for difference with the between-participant predictions was larger for the cognitive model (Cohen's d for tau 6.593 versus 5.195, and for MSE -6.803 versus -6.237).

#### III. Supplementary Figures

**Figure S1: Model validation.** A-B show that it is possible to differentiate within-individual use of different strategies and between-individual use of different strategies via a permutation test. Figures C-D show that the unsupervised (BMAPS-U) model inferences about strategy use when compared to out-of-scanner strategy use fall within the tolerance limits of within-individual variability in strategy use inferred from figures A-B, and outside the range of between-individual variability, thus validating the unsupervised inferences. Comparing supervised model inferences using SVMs with model (BMAPS-S) provide better estimates than using SVMs with behavioral and neuropsychological measures (behavioral-S). E-F show the effect size of differences between the models and between / within effects. Better model performance is indicated by a large effect size vs Between and a small effect size vs Within.

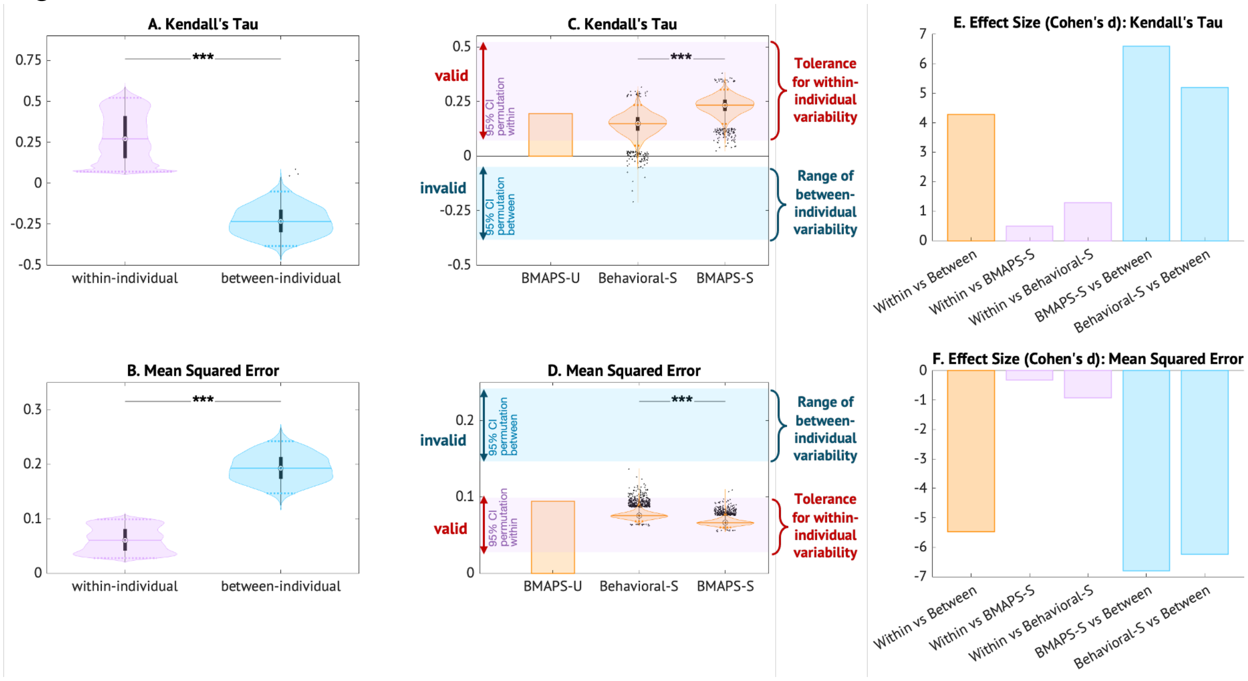

**Figure S2: Hierarchical clustering of items.**

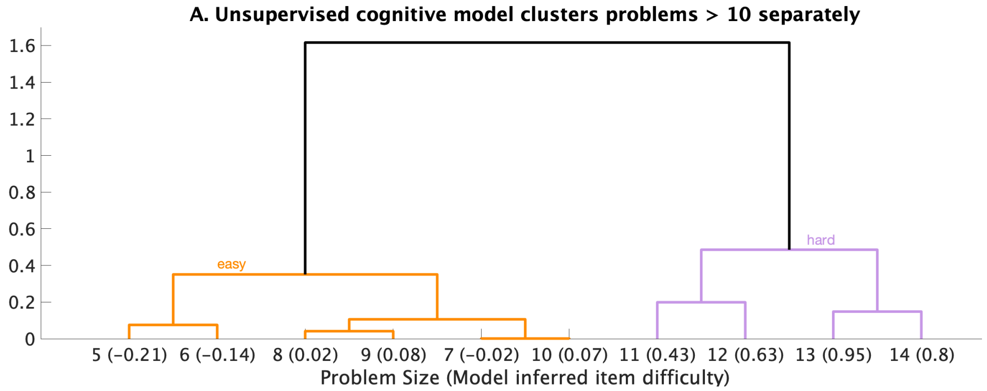

**Figure S3: Alternate SEM (control model) with affective measures (anxiety, attitude) modeled as outcomes of behavioral and cognitive measures.**

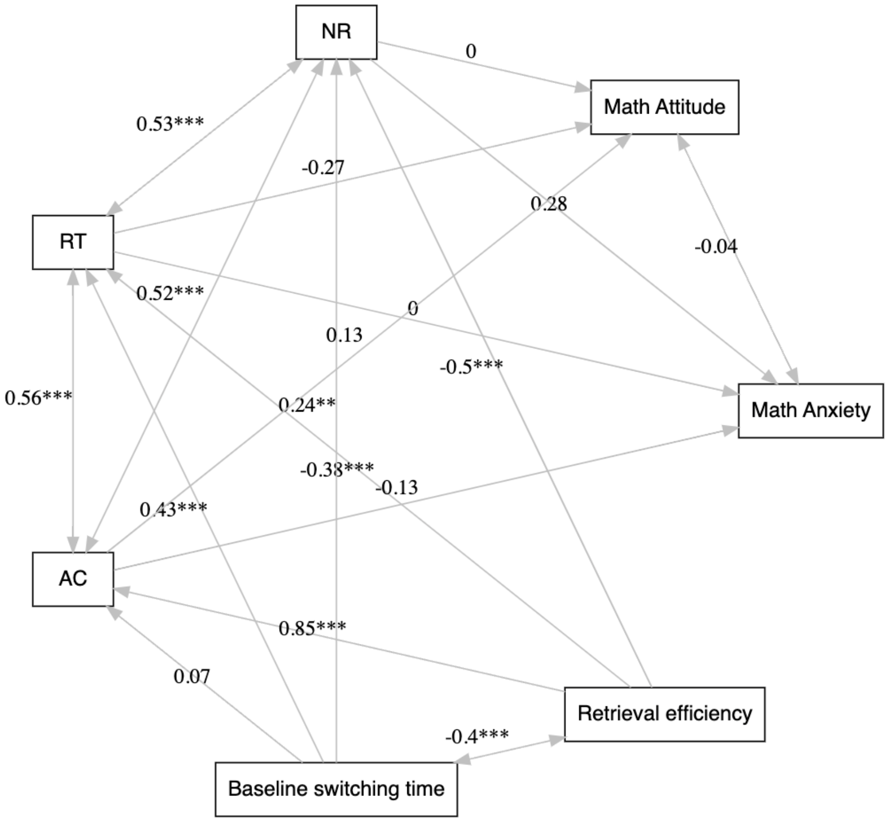

**Figure S4: Clusters in the BMAPS factor-space**

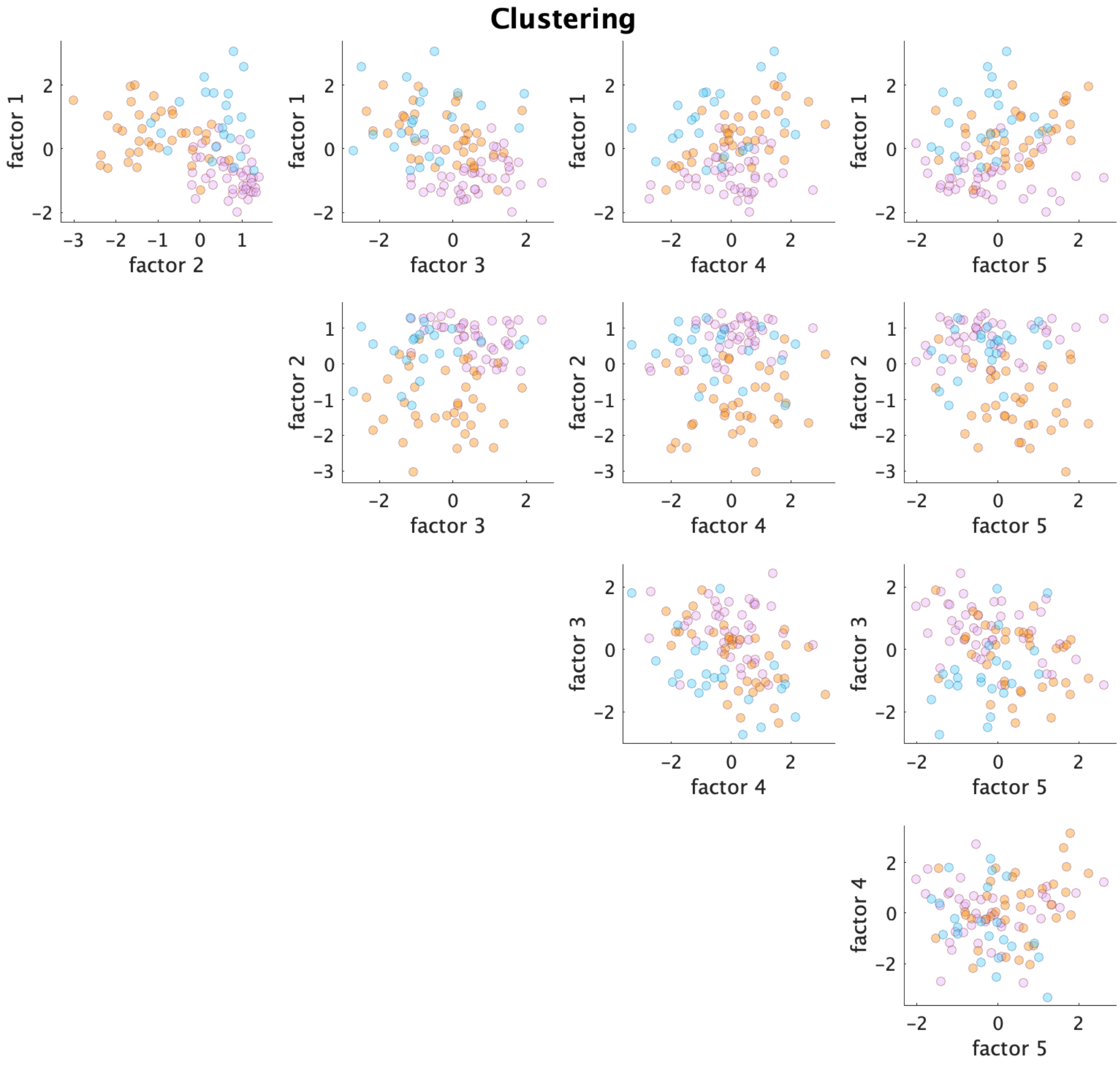

#### III. Supplementary Tables

**Supplementary Table 1:** Tolerance benchmarks based on within-individual and between-individual variability and comparison of model inferences about strategy use vs self-reports

| | **Kendal's tau** | | | **MSE** | | |
|---|---|---|---|---|---|---|
| | 95% CI | | Mean | 95% CI | | Mean |
| | Lower | Upper | | Lower | Upper | |
| Self-report within-individual | 0.071 | 0.522 | 0.283 | 0.028 | 0.099 | 0.062 |
| Self-report between-individual | -0.384 | 0.050 | -0.229 | 0.147 | 0.242 | 0.194 |
| BMAPS-Unsupervised vs self-report | | | 0.193 | | | 0.094 |
| Behavioral-Supervised vs self-report | | | 0.146 | | | 0.076 |
| BMAPS-Supervised vs self-report | | | 0.231 | | | 0.067 |

**Supplementary Table 2:** Significant correlations between model parameters. All other pairs of correlations between BMAPS measures were not significant (p > 0.05 after adjusting for multiple comparisons).

| Parameter 1 | Type | Parameter 2 | Type | r | p (unadjusted) | p (adjusted) |
|---|---|---|---|---|---|---|
| $\tau$ | C | $\alpha$ | C | -0.3909 | 0.000037 | 0.0041 |
| $\delta$ | C | $\tau$ | C | 0.4373 | 0.000003 | 0.0004 |
| $\beta$ | C | $\delta$ | C | 0.5128 | 0.000000 | 0.0000 |
| $\gamma$ | M | $\alpha$ | C | 0.4905 | 0.000000 | 0.0000 |
| $\gamma$ | M | $\tau$ | C | -0.4776 | 0.000000 | 0.0000 |
| $\gamma$ | M | $\delta$ | C | -0.4186 | 0.000010 | 0.0011 |
| $\Omega$ | M | $\mu$ | M | 0.5687 | 0.000000 | 0.0000 |
| $\gamma$ | M | $\Omega$ | M | -0.5502 | 0.000000 | 0.0000 |
| $\psi$ | M | $\Omega$ | M | 0.3749 | 0.000081 | 0.0090 |
| $\Sigma$ | M | $\Omega$ | M | -0.3724 | 0.000099 | 0.0108 |
| $\psi$ | M | $\gamma$ | M | -0.5123 | 0.000000 | 0.0000 |
| $\Sigma$ | M | $\gamma$ | M | 0.6079 | 0.000000 | 0.0000 |
| $\Sigma$ | M | $\psi$ | M | -0.4074 | 0.000018 | 0.0020 |

**Supplementary Table 3:** Correlations between BMAPS parameters and task behavior

| Parameter | Accuracy (AC) | | | Reaction Time (RT) | | | No Response (NR) | | |
|---|---|---|---|---|---|---|---|---|---|
| | **r** | **p** | **p (adj)** | **r** | **p** | **p (adj)** | **r** | **p** | **p (adj)** |
| $\alpha$ | 0.246 | 0.012 | 0.194 | **0.759** | **0.000** | **0.000** | **0.686** | **0.000** | **0.000** |
| $\kappa$ | 0.268 | 0.006 | 0.112 | -0.025 | 0.804 | 1.0 | -0.083 | 0.400 | 1.0 |
| $\tau$ | 0.189 | 0.054 | 0.652 | -0.165 | 0.092 | 0.918 | -0.269 | 0.006 | 0.111 |
| $\delta$ | **0.790** | **0.000** | **0.000** | **-0.551** | **0.000** | **0.000** | **-0.599** | **0.000** | **0.000** |
| $\beta$ | **0.443** | **0.000** | **0.000** | **-0.306** | **0.002** | **0.033** | **-0.305** | **0.002** | **0.033** |
| $\mu$ | 0.027 | 0.788 | 1.0 | **-0.380** | **0.000** | **0.002** | -0.111 | 0.261 | 1.0 |
| $\Omega$ | 0.016 | 0.873 | 1.0 | -0.256 | 0.008 | 0.151 | 0.026 | 0.789 | 1.0 |
| $\omega$ | 0.039 | 0.698 | 1.0 | 0.248 | 0.011 | 0.194 | **0.318** | **0.001** | **0.025** |
| $\gamma$ | -0.228 | 0.020 | 0.285 | **0.610** | **0.000** | **0.000** | **0.473** | **0.000** | **0.000** |
| $\psi$ | **0.382** | **0.000** | **0.002** | -0.126 | 0.201 | 1.0 | -0.048 | 0.626 | 1.0 |
| $\Sigma$ | -0.231 | 0.019 | 0.285 | 0.217 | 0.027 | 0.346 | 0.173 | 0.079 | 0.866 |

**Supplementary Table 4:** Correlations between BMAPS derived factors and task behavior

| Factor | Accuracy (AC) | | | Reaction Time (RT) | | | No Response (NR) | | |
|---|---|---|---|---|---|---|---|---|---|
| | r | p | p (adj) | r | p | p (adj) | r | p | p (adj) |
| 1 | **-0.334** | **0.001** | **0.006** | **0.399** | **0.000** | **0.000** | **0.327** | **0.001** | **0.006** |
| 2 | 0.043 | 0.671 | 1.0 | **-0.353** | **0.000** | **0.003** | -0.026 | 0.794 | 1.0 |
| 3 | **0.621** | **0.000** | **0.000** | **-0.430** | **0.000** | **0.000** | **-0.441** | **0.000** | **0.000** |
| 4 | 0.004 | 0.972 | 1.0 | -0.013 | 0.897 | 1.0 | -0.007 | 0.942 | 1.0 |
| 5 | **0.287** | **0.004** | **0.023** | **0.767** | **0.000** | **0.000** | **0.675** | **0.000** | **0.000** |

**Supplementary Table 5: Sex differences**

| Parameter | t-stat | df | p | p (adj) |
|---|---|---|---|---|
| $\alpha$ | 0.181 | 103 | 0.857 | 1.000 |
| $\kappa$ | -2.362 | 103 | 0.020 | 0.321 |
| $\tau$ | -0.906 | 103 | 0.367 | 1.000 |
| $\delta$ | -0.044 | 102 | 0.965 | 1.000 |
| $\beta$ | -1.248 | 103 | 0.215 | 1.000 |
| $\mu$ | -0.765 | 103 | 0.446 | 1.000 |
| Ω | -0.805 | 103 | 0.423 | 1.000 |
| $\omega$ | 1.555 | 101 | 0.123 | 1.000 |
| $\gamma$ | 1.363 | 103 | 0.176 | 1.000 |
| ψ | -1.918 | 103 | 0.058 | 0.868 |
| Σ | 0.466 | 102 | 0.642 | 1.000 |
| Factor 1 | 1.391 | 99 | 0.167 | 1.000 |
| Factor 2 | -1.039 | 99 | 0.302 | 1.000 |
| Factor 3 | -1.560 | 99 | 0.122 | 1.000 |
| Factor 4 | -0.885 | 99 | 0.378 | 1.000 |
| Factor 5 | 0.126 | 99 | 0.900 | 1.000 |

**Supplementary Table 6: Correlations between BMAPS parameters and WIAT-II measures of math achievement**

| BMAPS parameter | | WIAT-II | r | p (unadjusted) | p (adjusted) |
|---|---|---|---|---|---|
| mean entropy | $\Omega$ | *NumOps* | 0.328 | 0.00088 | 0.0071 |
| baseline switch time | $\gamma$ | *NumOps* | -0.413 | 0.00002 | 0.0002 |
| adaptivity | $\psi$ | *NumOps* | 0.343 | 0.00048 | 0.0043 |
| SD entropy | $\Sigma$ | *NumOps* | -0.393 | 0.00006 | 0.0006 |
| retrieval efficiency | $\delta$ | *MathReas* | 0.318 | 0.0011 | 0.0102 |
| baseline switch time | $\gamma$ | *MathReas* | -0.393 | < 0.00001 | 0.0004 |
| adaptivity | $\psi$ | *MathReas* | 0.336 | 0.0005 | 0.0053 |
| SD entropy | $\Sigma$ | *MathReas* | -0.281 | 0.0042 | 0.0337 |

**Supplementary Table 7: Stepwise linear regression results – Numerical Operations**

| | Estimate | SE | t-stat | p - value |
|---|---|---|---|---|
| Intercept | -1.4e-16 | 0.092 | -1.5e-15 | 1 |
| $\gamma$ | - 0.516 | 0.104 | - 4.96 | 3.16e-6 |
| $\tau$ | - 0.216 | 0.104 | - 2.07 | 0.0409 |
| N (observations) | 96 | | | |
| Error df | 93 | | | |
| F-stat vs Constant | 12.3 | | | |
| p - value | 1.77e-05 | | | |
| Ordinary $R^2$ | 0.2097 | | | |
| Adjusted $R^2$ | 0.1927 | | | |

**Supplementary Table 8: Stepwise linear regression results – Mathematical Reasoning**

| | **Estimate** | **SE** | **t-stat** | **p - value** |
|---|---|---|---|---|
| Intercept | 3.74e-17 | 0.0089 | 4.19e-16 | 1 |
| $\gamma$ | -0.412 | 0.113 | -3.65 | 0.0004 |
| $\delta$ | 0.229 | 0.099 | 2.33 | 0.0222 |
| $\alpha$ | 0.228 | 0.105 | 2.18 | 0.0315 |
| N (observations) | 99 | | | |
| Error df | 95 | | | |
| F-stat vs Constant | 9.81 | | | |
| p - value | 1.07e-05 | | | |
| Ordinary $R^2$ | 0.2365 | | | |
| Adjusted $R^2$ | 0.2124 | | | |

#### Supplementary Table 9: Exploratory and control models for mediation effects

| | | $\mathbf{p\ (\chi^2)}$ | CFI | TLI | RMSEA | $\boldsymbol{\beta_{XY}}$ | $\mathbf{p\ (\beta_{XY})}$ |
|---|---|---|---|---|---|---|---|
| **Threshold** | | **>= 0.05** | **>= 0.95** | **>= 0.95** | **<= 0.08** | | **<= 0.05** |
| $\alpha$ | r-ATM | **0.055** | **0.979** | 0.931 | 0.129 | -0.162 | 0.115 |
| $\boldsymbol{\delta}$ | **r-ATM** | **0.344** | **0.999** | **0.995** | **0.035** | **0.279** | **0.006** |
| $\tau$ | r-ATM | 0.017 | 0.914 | 0.713 | 0.161 | 0.023 | 0.825 |
| $\beta$ | r-ATM | **0.082** | **0.966** | 0.886 | 0.116 | 0.203 | **0.046** |
| $\kappa$ | r-ATM | 0.024 | 0.923 | 0.745 | 0.152 | 0.098 | 0.346 |
| $\mu$ | r-ATM | 0.016 | 0.922 | 0.741 | 0.164 | 0.041 | 0.696 |
| $\omega$ | r-ATM | 0.049 | 0.946 | 0.819 | 0.134 | -0.205 | 0.047 |
| $\Omega$ | r-ATM | 0.028 | 0.934 | 0.779 | 0.149 | 0.107 | 0.302 |
| $\boldsymbol{\gamma}$ | **r-ATM** | **0.362** | **0.998** | **0.995** | **0.027** | **-0.303** | **0.002** |
| $\Psi$ | r-ATM | 0.017 | 0.927 | 0.756 | 0.161 | 0.047 | 0.649 |
| $\Sigma$ | r-ATM | **0.081** | **0.953** | 0.843 | 0.117 | -0.169 | 0.101 |
| $\alpha$ | sema-C | 0.020 | **0.972** | 0.906 | 0.152 | 0.129 | 0.197 |
| $\boldsymbol{\delta}$ | **sema-C** | **0.411** | **1.000** | **1.002** | **0.000** | **-0.278** | **0.004** |
| $\tau$ | sema-C | 0.011 | 0.911 | 0.702 | 0.166 | -0.061 | 0.546 |
| $\beta$ | sema-C | 0.032 | 0.947 | 0.823 | 0.140 | -0.160 | 0.106 |
| $\kappa$ | sema-C | 0.014 | 0.913 | 0.711 | 0.160 | -0.102 | 0.307 |
| $\mu$ | sema-C | 0.009 | 0.918 | 0.726 | 0.170 | 0.008 | 0.937 |
| $\omega$ | sema-C | 0.013 | 0.914 | 0.714 | 0.164 | 0.055 | 0.585 |
| $\Omega$ | sema-C | 0.010 | 0.916 | 0.719 | 0.168 | -0.027 | 0.788 |
| $\gamma$ | sema-C | **0.055** | **0.964** | 0.881 | 0.124 | 0.202 | **0.040** |
| $\Psi$ | sema-C | 0.012 | 0.918 | 0.727 | 0.164 | -0.056 | 0.575 |
| $\Sigma$ | sema-C | 0.037 | 0.939 | 0.796 | 0.137 | 0.247 | **0.012** |

**Supplementary Table 10: Longitudinal changes in parameters**

| Parameter | t-stat | df | p | p (adj) |
|---|---|---|---|---|
| $\alpha$ | -4.76 | 29 | $< 0.0001$ | 0.0005 |
| $\delta$ | 3.97 | 29 | 0.0004 | 0.0039 |
| $\gamma$ | -4.49 | 29 | 0.0001 | 0.0011 |

**Supplementary Table 11: Stepwise linear regression results – Mathematical Reasoning at second time point**

| | **Estimate** | **SE** | **t-stat** | **p - value** |
|---|---|---|---|---|
| Intercept | 7.9e-16 | 0.133 | -0.419 | 0.679 |
| Year 2 $\gamma$ | -0.404 | 0.136 | 2.249 | 0.033 |
| N (observations) | 29 | | | |
| Error df | 26 | | | |
| F-stat vs Constant | 14.3 | | | |
| p - value | 6.36e-05 | | | |
| Ordinary $R^2$ | 0.525 | | | |
| Adjusted $R^2$ | 0.488 | | | |

**Supplementary Table 12: Longitudinal ICC**

| Consistency | ICC | LB | UB | F | df1 | df2 | p | Adj. p |
|---|---|---|---|---|---|---|---|---|
| $\alpha$ | 0.259 | -0.106 | 0.562 | 1.698 | 29.000 | 29.000 | 0.080 | 0.640 |
| $\delta$ | 0.315 | -0.045 | 0.603 | 1.920 | 29.000 | 29.000 | 0.042 | 0.379 |
| $\tau$ | -0.123 | -0.458 | 0.243 | 0.781 | 29.000 | 29.000 | 0.745 | 0.752 |
| $\beta$ | 0.059 | -0.302 | 0.406 | 1.126 | 29.000 | 29.000 | 0.376 | 0.891 |
| $\kappa$ | 0.362 | 0.008 | 0.635 | 2.135 | 29.000 | 29.000 | 0.023 | 0.250 |
| $\mu$ | 0.106 | -0.258 | 0.445 | 1.238 | 29.000 | 29.000 | 0.284 | 1.000 |
| $\omega$ | 0.390 | 0.034 | 0.658 | 2.279 | 28.000 | 28.000 | 0.017 | 0.199 |
| $\Omega$ | 0.100 | -0.265 | 0.439 | 1.221 | 29.000 | 29.000 | 0.297 | 1.000 |
| $\gamma$ | 0.337 | -0.021 | 0.618 | 2.015 | 29.000 | 29.000 | 0.032 | 0.320 |
| $\psi$ | 0.127 | -0.245 | 0.467 | 1.291 | 28.000 | 28.000 | 0.252 | 1.000 |
| $\Sigma$ | 0.115 | -0.251 | 0.451 | 1.259 | 29.000 | 29.000 | 0.270 | 1.000 |
| **RT** | 0.454 | 0.118 | 0.697 | 2.664 | 29.000 | 29.000 | 0.005 | 0.072 |
| **AC** | 0.447 | 0.095 | 0.699 | 2.616 | 27.000 | 27.000 | 0.008 | 0.098 |
| **NR** | 0.187 | -0.193 | 0.519 | 1.461 | 27.000 | 27.000 | 0.165 | 1.000 |
| **Agreement** | **ICC** | **LB** | **UB** | **F** | **df1** | **df2** | **p** | **Adj. p** |
| $\alpha$ | 0.168 | -0.098 | 0.450 | 1.698 | 29.000 | 17.954 | 0.121 | 0.971 |
| $\delta$ | 0.236 | -0.071 | 0.524 | 1.920 | 29.000 | 19.109 | 0.070 | 0.760 |
| $\tau$ | -0.127 | -0.476 | 0.248 | 0.781 | 29.000 | 28.866 | 0.745 | 0.750 |
| $\beta$ | 0.057 | -0.283 | 0.394 | 1.126 | 29.000 | 29.442 | 0.375 | 0.890 |
| $\kappa$ | 0.363 | 0.011 | 0.635 | 2.135 | 29.000 | 29.714 | 0.022 | 0.283 |
| $\mu$ | 0.098 | -0.228 | 0.421 | 1.238 | 29.000 | 29.889 | 0.282 | 1.000 |
| $\omega$ | 0.342 | 0.008 | 0.617 | 2.279 | 28.000 | 23.752 | 0.022 | 0.283 |
| $\Omega$ | 0.101 | -0.270 | 0.443 | 1.221 | 29.000 | 29.161 | 0.297 | 1.000 |
| $\gamma$ | 0.237 | -0.075 | 0.527 | 2.015 | 29.000 | 15.544 | 0.074 | 0.701 |
| $\psi$ | 0.128 | -0.246 | 0.468 | 1.291 | 28.000 | 28.314 | 0.251 | 1.000 |
| $\Sigma$ | 0.118 | -0.260 | 0.459 | 1.259 | 29.000 | 29.043 | 0.270 | 1.000 |
| **RT** | 0.296 | -0.078 | 0.603 | 2.664 | 29.000 | 8.438 | 0.069 | 0.760 |
| **AC** | 0.418 | 0.081 | 0.676 | 2.616 | 27.000 | 25.937 | 0.008 | 0.116 |
| **NR** | 0.101 | -0.108 | 0.362 | 1.461 | 27.000 | 18.630 | 0.200 | 1.000 |

#### Supplementary Table 13: Clustering ANOVA results – baseline measures

| Measure | F-stat | p | Adj p | Mean Values | | |
|---|---|---|---|---|---|---|
| | | | | Cluster 1 "good" | Cluster 2 "perfectionist" | Cluster 3 "not-so-good" |
| **$\alpha$** | **5.24** | **0.00690** | **0.04140** | **3.78** | **4.21** | **3.73** |
| **$\delta$** | **19.39** | **0.00000** | **0.00000** | **1.71** | **1.40** | **1.12** |
| $\tau$ | 4.75 | 0.01070 | 0.05360 | 0.74 | 0.53 | 0.68 |
| **$\beta$** | **5.79** | **0.00420** | **0.03360** | **0.55** | **0.53** | **0.52** |
| **$\kappa$** | **10.47** | **0.00010** | **0.00080** | **3.13** | **3.18** | **2.72** |
| **$\mu$** | **42.45** | **0.00000** | **0.00000** | **-0.07** | **-1.96** | **-0.41** |
| **$\omega$** | **10.27** | **0.00010** | **0.00080** | **0.40** | **0.67** | **0.47** |
| **$\Omega$** | **52.71** | **0.00000** | **0.00000** | **0.65** | **0.49** | **0.62** |
| **$\gamma$** | **37.41** | **0.00000** | **0.00000** | **0.53** | **1.03** | **1.02** |
| **$\psi$** | **28.77** | **0.00000** | **0.00000** | **0.49** | **0.30** | **0.22** |
| **$\Sigma$** | **31.36** | **0.00000** | **0.00000** | **0.13** | **0.17** | **0.20** |
| **MR (WIAT)** | **10.85** | **0.00010** | **0.00070** | **115.24** | **107.19** | **100.14** |
| **NO (WIAT)** | **10.40** | **0.00010** | **0.00080** | **111.33** | **100.18** | **97.41** |
| SEMA-C | 2.40 | 0.09610 | 0.19230 | 5.10 | 5.59 | 8.00 |
| **RT** | **12.81** | **0.00000** | **0.00010** | **2.33** | **2.82** | **2.71** |
| **AC** | **24.84** | **0.00000** | **0.00000** | **0.94** | **0.91** | **0.83** |
| NR | 4.47 | 0.01390 | 0.05570 | 0.08 | 0.12 | 0.14 |
| **Factor 1** | **44.40** | **0.00000** | **0.00000** | **-0.84** | **0.43** | **0.88** |
| **Factor 2** | **69.97** | **0.00000** | **0.00000** | **0.73** | **-1.07** | **0.40** |
| **Factor 3** | **14.62** | **0.00000** | **0.00000** | **0.60** | **-0.24** | **-0.74** |
| Factor 4 | 2.12 | 0.12540 | 0.19230 | 0.02 | 0.24 | -0.45 |
| **Factor 5** | **5.68** | **0.00470** | **0.03360** | **-0.22** | **0.42** | **-0.30** |
| Math attitude | 4.07 | 0.02050 | 0.06140 | 1.14 | 0.89 | 0.84 |
| **Retrieval %** | **35.00** | **0.00000** | **0.00000** | **0.44** | **0.15** | **0.38** |

#### Supplementary Table 14: Clustering ANOVA results – longitudinal changes

| Measure | F-stat | p | Adj p | Mean Values | | |
|---|---|---|---|---|---|---|
| | | | | **Cluster 1** | **Cluster 2** | **Cluster 3** |
| $\boldsymbol{\alpha}$ | 0.0895 | 0.9147 | 1.7892 | -0.5992 | -0.4921 | -0.5239 |
| $\boldsymbol{\delta}$ | 0.1119 | 0.8946 | 1.7892 | 0.5532 | 0.4254 | 0.5491 |
| $\tau$ | 1.5735 | 0.2264 | 1.1775 | -0.0454 | 0.2252 | -0.1051 |
| $\boldsymbol{\beta}$ | 0.7655 | 0.4753 | 1.4570 | 0.0029 | 0.0095 | 0.0261 |
| $\boldsymbol{\kappa}$ | 3.3412 | 0.0511 | 0.4089 | 0.2092 | -0.4417 | -0.0909 |
| $\boldsymbol{\mu}$ | **6.5524** | **0.0050** | **0.0496** | **0.0081** | **1.4705** | **-0.5331** |
| $\boldsymbol{\omega}$ | 1.0502 | 0.3642 | 1.4570 | -0.1362 | -0.0570 | -0.2379 |
| $\boldsymbol{\Omega}$ | **11.3982** | **0.0003** | **0.0031** | **-0.0341** | **0.0932** | **-0.0692** |
| $\boldsymbol{\gamma}$ | 1.7348 | 0.1962 | 1.1775 | -0.1438 | -0.3287 | -0.1956 |
| $\boldsymbol{\psi}$ | 2.4244 | 0.1091 | 0.7634 | -0.0489 | 0.1625 | 0.0381 |
| $\boldsymbol{\Sigma}$ | 4.7402 | 0.0176 | 0.1581 | 0.0233 | -0.0235 | -0.0236 |
